# Whole genome sequencing of *Borrelia burgdorferi* isolates reveals linked clusters of plasmid-borne accessory genome elements associated with virulence

**DOI:** 10.1101/2023.02.26.530159

**Authors:** Jacob E. Lemieux, Weihua Huang, Nathan Hill, Tjasa Cerar, Lisa Freimark, Sergio Hernandez, Matteo Luban, Vera Maraspin, Petra Bogovic, Katarina Ogrinc, Eva Ruzic-Sabljic, Pascal Lapierre, Erica Lasek-Nesselquist, Navjot Singh, Radha Iyer, Dionysios Liveris, Kurt D. Reed, John M. Leong, John A. Branda, Allen C. Steere, Gary P. Wormser, Franc Strle, Pardis C. Sabeti, Ira Schwartz, Klemen Strle

## Abstract

Lyme disease is the most common vector-borne disease in North America and Europe. The clinical manifestations of Lyme disease vary based on the genospecies of the infecting *Borrelia burgdorferi* spirochete, but the microbial genetic elements underlying these associations are not known. Here, we report the whole genome sequence (WGS) and analysis of 299 patient-derived *B. burgdorferi* sensu stricto (*Bbss*) isolates from patients in the Eastern and Midwestern US and Central Europe. We develop a WGS-based classification of *Bbss* isolates, confirm and extend the findings of previous single- and multi-locus typing systems, define the plasmid profiles of human-infectious *Bbss* isolates, annotate the core and strain-variable surface lipoproteome, and identify loci associated with disseminated infection. A core genome consisting of ∼800 open reading frames and a core set of plasmids consisting of lp17, lp25, lp36, lp28-3, lp28-4, lp54, and cp26 are found in nearly all isolates. Strain-variable (accessory) plasmids and genes correlate strongly with phylogeny. Using genetic association study methods, we identify an accessory genome signature associated with dissemination and define the individual plasmids and genes that make up this signature. Strains within the RST1/WGS A subgroup, particularly a subset marked by the OspC type A genotype, are associated with increased rates of dissemination. OspC type A strains possess a unique constellation of strongly linked genetic changes including the presence of lp56 and lp28-1 plasmids and a cluster of genes that may contribute to their enhanced virulence compared to other genotypes. The patterns of OspC type A strains typify a broader paradigm across *Bbss* isolates, in which genetic structure is defined by correlated groups of strain-variable genes located predominantly on plasmids, particularly for expression of surface-exposed lipoproteins. These clusters of genes are inherited in blocks through strain-specific patterns of plasmid occupancy and are associated with the probability of invasive infection.

## INTRODUCTION

Lyme disease is a heterogeneous illness caused by spirochetes of the *Borrelia burgdorferi* sensu lato (*Bbsl*, sensu lato meaning ‘in the broad sense’) complex. *Bbsl* contains over 20 subspecies (also termed genospecies, genomic species), four of which cause the majority of disease in humans: *B. burgdorferi* sensu stricto (*Bbss,* sensu stricto meaning in the strict sense), *B. afzelii*, *B. garinii*, and *B. bavariensis* [1]. Nearly all Lyme disease in the US is caused by *Bbss*. In Europe, most infections are caused by *B. afzelii*, *B. garinii*, or *B. bavariensis,* whereas infection due to *Bbss* is rare. Infection with *Bbsl* usually presents as an expanding skin rash, erythema migrans (EM), at the site of the tick-bite. If untreated, spirochetes may disseminate to secondary sites (a phenotype described as ‘dissemination’), primarily other skin sites, the nervous system and joints [1, 2]. In addition to clinical variation among *Bbsl* species, differences in virulence have also been noted between genotypes within *Bbss* [3–5], and such phenotypes have been recapitulated in murine models [6–8]. These associations imply that microbial genetic loci likely influence the clinical manifestations of Lyme disease. Despite such evidence linking microbial genotype to clinical phenotype, the specific genes or loci responsible for the clinical manifestations of Lyme disease have not yet been identified.

*Bbss* genome analysis has been limited to date due to technical challenges of sequencing and assembly and difficulties of obtaining isolates from cases of human disease. The *Bbss* genome consists of a roughly one megabase of core genome (consisting of a ∼900Kb chromosome and the plasmids cp26 and lp54), as well as numerous (>15) additional circular and linear extrachromosomal DNA elements (colloquially termed plasmids) [9, 10]. Subsets of plasmids have high levels of homology (as exemplified by seven 32 kilobase circular plasmids (cp32) [11] and four 28-kilobase linear plasmids (lp28) [10] in the B31 reference isolate), which have diversified through duplication, recombination, and other primordial evolutionary events [12], The sheer number of plasmids and their extreme homology has made sequencing and assembly of complete *Bbss* genomes a major challenge, particularly with widely-used short read sequencing methods [13].

The technical challenges of sequencing and assembly are compounded by the difficulty of obtaining isolates from human disease. It has been possible to culture the organism from EM lesions in the majority of cases, but this requires a skin biopsy and specialized culture techniques, both of which are rarely used in routine clinical practice. The organism has occasionally been cultured from CSF in patients with meningitis, but extremely rarely from synovial fluid in patients with Lyme arthritis, the most common late disease manifestation in the US. Thus, the great majority of available *Bbss* isolates are from patients with EM, an early disease manifestation. As a result of these challenges, only a small number of human clinical isolates have been sequenced and analyzed. To our knowledge, no large WGS studies of human isolates have been conducted. Fewer than 50 human isolates analyzed by WGS have been publicly reported, either sporadically or included in cohorts consisting primarily of tick-derived isolates [14–19].

Genotyping systems have been developed to subclassify *Bbss* strains using single or multiple genomic regions (reviewed in [20]). Two of the most commonly used typing methods are based on a restriction-fragment length polymorphisms in the 16S-23S ribosomal RNA spacer region [21, 22], termed ribosomal spacer type (RST), and on sequence variation of outer surface protein C (OspC), one of the most variable *Bbss* proteins [23, 24]. RST typing subdivides *Bbss* into 3 types, referred to as RST1, RST2, and RST3 [6], whereas OspC typing subdivides *Bbss* into ∼30 OspC genotypes of which >24 cause infection in humans [25–27]. RST and OspC are in linkage disequilibrium on the core genome, and each RST genotype is generally associated with particular OspC types (e.g., RST1 mostly corresponds to OspC types A and B and RST2 corresponds primarily to OspC types F, H, K and N) [27]), whereas RST3 is the most variable and correlates with the remaining OspC types. In addition to these genotyping methods, multilocus sequence typing (MLST), which is based on eight chromosomal housekeeping genes, has been used to further sub-stratify the strains [27, 28]. According to the *Borrelia* MLST database (https://pubmlst.org/borrelia/), >900 MLST sequence types have been identified.

Application of targeted genotyping methods has previously established a link between *Bbss* microbial genotype and several phenotypic properties including dissemination, disease severity, immunogenicity, and distinct clinical presentation [1,3,5,6,8,26,27,29–32]. For example, using RST and OspC genotyping we previously showed that RST1 OspC type A strains have greater proclivity to disseminate, are more immunogenic, are associated with more symptomatic early infection, and with a greater frequency of post-infectious Lyme arthritis. However, these approaches lack the resolution to reconstruct a detailed evolutionary history or to define individual genes or loci underlying phenotypic variability. The limitations of previous studies have been further compounded by the absence of large cohorts of patient-derived isolates accompanied by detailed clinical information. Here, we used whole genome sequencing to characterize in detail the genomes – including the core genome and associated plasmids – of 299 patient-derived *Bbss* strains. The isolates were collected primarily from patients with EM, the initial skin lesion of the infection, over three decades across Northeastern and Midwestern US and Central Europe. We carried out phylogenetic and phylogeographic analysis, and identified particular *Bbss* genomic groups, plasmids, and individual open reading frames (ORFs) associated with tissue invasive (disseminated) human disease.

## MATERIALS and METHODS

### Selection of *B. burgdorferi* isolates (see Supplemental Table 1)

In total, 299 *Bbss* isolates collected from 299 patients over a 30-year period (1992-2021) were included in this study: 202 from the Northeastern US, 61 from the Midwestern US and 36 from Slovenia (Central Europe). The majority (97%) of isolates were derived from skin (n = 287) or blood (n = 2) of patients (9 were derived from cerebrospinal fluid [CSF]) by culturing in BSK or MKP medium [33, 34]. All patients met the US Centers for Disease Control and Prevention (CDC) criteria for Lyme disease [35]. Only low passage isolates (passage <5) were used for WGS.

#### Northeastern United States

The 201 isolates from the Northeastern US were collected at two geographic locations: 113 from New England (primarily from contiguous regions of Massachusetts, Rhode Island, and Connecticut) and 88 from New York State. The New York strains belong to a larger collection of more than 400 clinical isolates, collected between 1992-2005, that had been previously typed at the *rrs-rrlA* IGS and *ospC* loci [4, 31]. To account for the full diversity of *Bbss* genotypes found in the collection, isolates with the best sequence quality from each OspC major group were selected for this study in accordance with their prevalence in the entire collection. All of the latter isolates were cultured from skin biopsies of infected patients, rather than from blood or CSF (Supplemental Tables 1 and 2).

#### Midwestern United States

The 62 isolates from the Midwestern US were derived from specimens submitted to the Marshfield Laboratories (Marshfield, WI) for *Borrelia* culture from 1993 to 2003 (Supplemental Tables 1 and 2).

#### Central Europe (Slovenia)

The 36 isolates from Slovenia represent all *Bbss* isolates that were cultured from patients over a 27-year period (1994-2021), who were evaluated at the Lyme borreliosis outpatient clinic at the University Medical Center Ljubljana (UMCL).

### Selection of patients

This study involves secondary use of deidentified archival clinical isolates and patient data collected in previous studies and was approved by the Massachusetts General Hospital Institutional Review Board (IRB) under protocol 2019P001864. Patients included in this study were diagnosed with early Lyme disease and were classified as having either localized or disseminated infection. Early Lyme disease was defined by the presence of at least one EM skin lesion or symptoms consistent with Lyme neuroborreliosis along with a positive CSF culture. Localized infection was defined by a single culture positive EM skin lesion in the absence of clinical and/or microbiological evidence of dissemination to a secondary site. Disseminated infection was defined by a positive blood or CSF culture or PCR, multiple EM lesions, and/or signs of neurological involvement. We were able to classify 291 or the 299 (97.3%) isolates as Disseminated or Localized by these criteria. Clinical records were not available to classify 8/299 (2.7%), and these isolates were excluded from analyses of dissemination. A measure of bloodstream dissemination was available for 212/299 (70.9%) of isolates, with blood PCR available for 106/299 (35.4%) and blood culture available for a disjoint set of 106/299 (35.4%) of all isolates. Multiple EM was present in 57 / 290 (19.7%); among patients with a single EM, 23/88 (26.1%) had a positive blood culture and 28/86 (32.6%) had a positive PCR. Complications such as Lyme neuroborreliosis were defined by clinical criteria and based on assessment by the treating clinician. In Europe, central nervous system (CNS) pleocytosis and intrathecal production of *Borrelia* antibodies were required for diagnostic determination of Lyme neuroborreliosis, following the EFNS guidelines [36]. Summary statistics of isolates by group is provided in Supplemental Table 1. The list of isolates and associated metadata is provided in Supplemental Table 2.

### Whole-Genome Sequencing

*Bbss* DNA was isolated from the cultured isolates with either the IsoQuick kit (Orca Research, Bothell, WA), the Gentra PureGene DNA Isolation Kit (Qiagen Inc., Valencia, CA), or the DNEasy kit (Qiagen Inc, Valencia, CA). Short-read next-generation sequencing (NGS) library construction was performed using the Nextera XT Library Prep Kit (Illumina, San Diego, CA). DNA quantification was performed in a 96-well microplate using the SpectraMax Quant dsDNA Assay Kit and the Gemini XPS Fluorometer (Molecular Devices, San Jose, CA), or in a single tube using the Qubit 2.0 fluorometer (Thermo Fisher Scientific, Springfield Township, NJ). Library quality was examined using the 4200 TapeStation and D1000 ScreenTape (Agilent, Santa Clara, CA). Paired-end sequencing (2 × 150 or 250 cycles) was performed using the NextSeq 550 or MiSeq system (Illumina).

### Bioinformatics Data Analysis

Trimmomatic v0.39 [37] was used for trimming and cleaning of raw sequence reads; SPAdes v3.14.1 [38] for *de novo* genome assembly; QUAST [39] for quality assessment and assembly visualization; Kraken2 [40] v2.1.1 for digital cleaning of assembled genomic sequence by using taxonomy classification; mlst v2.19.0 (https://github.com/tseemann) for MLST [41] identification from assembled sequences; k-mer weighted inner product (kWIP) [42] v0.2.0 for alignment-free, k-mer-based relatedness analysis; prokka v1.14.6 [43] for genome sequence annotation; Roary [44] for core- and pan-genome analysis; FastTree v2.1.11 [45] for phylogeny tree generation. Bioconductor [46] packages in R [47] v4.1.1 and/or RStudio v2021.09.0+351, such as ggplot2 [48], ggtree [49], ggtreeExtra, and ggstar, were also used for phylogeny tree generation. MLST definitions were downloaded from pubMLST. Multidimensional scaling (MDS) was calculated on the kWIP distances using the command mdscale() in R. Fisher’s exact test was used for pairwise comparison of categorical variables using the fisher.test() function in R. The MiniKraken2 database was constructed for Kraken2 from complete bacterial, archaeal, and viral genomes in RefSeq as of March 12, 2020. To characterize the plasmid content of individual isolates, we took two approaches. We first aligned the contigs to the B31 reference and quantified a plasmid as present or absent if greater than 50% of the reference genome plasmid was covered by contigs. As a complementary approach, we built a hidden Markov model (HMM) of PFam32 genes using HMMer [50] and searched the resulting profile against the assemblies to identify PFam32 genes. We then aligned the resulting putative PFam32 genes against a set of canonical PFam32 genes, provided by Dr. Sherwood Casjens, that have been used to determine plasmid types in published reports [51]. For each putative PFam32 gene, if a match with <5% identity was present in the list of annotated PFam32 genes, we marked the isolate as having a copy of the closest-matching PFam32 based on sequence identity. If no PFam32 within these thresholds could be identified, the closest PFam32 family member was considered unknown and not assigned in this analysis.

## RESULTS

### Whole-genome sequencing of human Borrelia burgdorferi sensu stricto isolates

To gain insight into the evolution, population structure, and pathogenesis of *Bbss* in human infection, we sequenced the complete genomes of 299 *Bbss* from human cases of early Lyme disease. We sequenced their whole genomes at a median coverage of 57.6x (interquartile range [IQR] 27.6x - 130.8x). The *de novo* assemblies produced high-quality, nearly-complete genomic assemblies with a median total length of 1.34 megabases (Mb) (IQR 1.30 - 1.37 Mb). Final assemblies contained a median of 107 contigs per isolate (IQR 88.0 - 137.5) and had a median N50 of 213,476 bases (IQR 80,809 - 221,506 bases). We were unable to finish assembly of plasmids due to repetitive plasmid sequences. Assembly statistics are given in Supplemental Table 3.

As an initial characterization of divergence between strains without any reference or annotation, we applied alignment-free, kmer-based analysis (kWIP) to the WGS data and identified three major clusters based on their genetic distances (Figure 1C and D, Figure S1). This unbiased distance analysis revealed that a single lineage (WGS A) was divergent from all other isolates (Figure 1C and D). The remaining isolates are grouped into two stable clusters (WGS groups B and C). RST type 1 was divergent from the other two WGS groups, but RST 2 and 3 were mixed between WGS groups B and C (Figures 1C and 1D).

**Figure 1:**
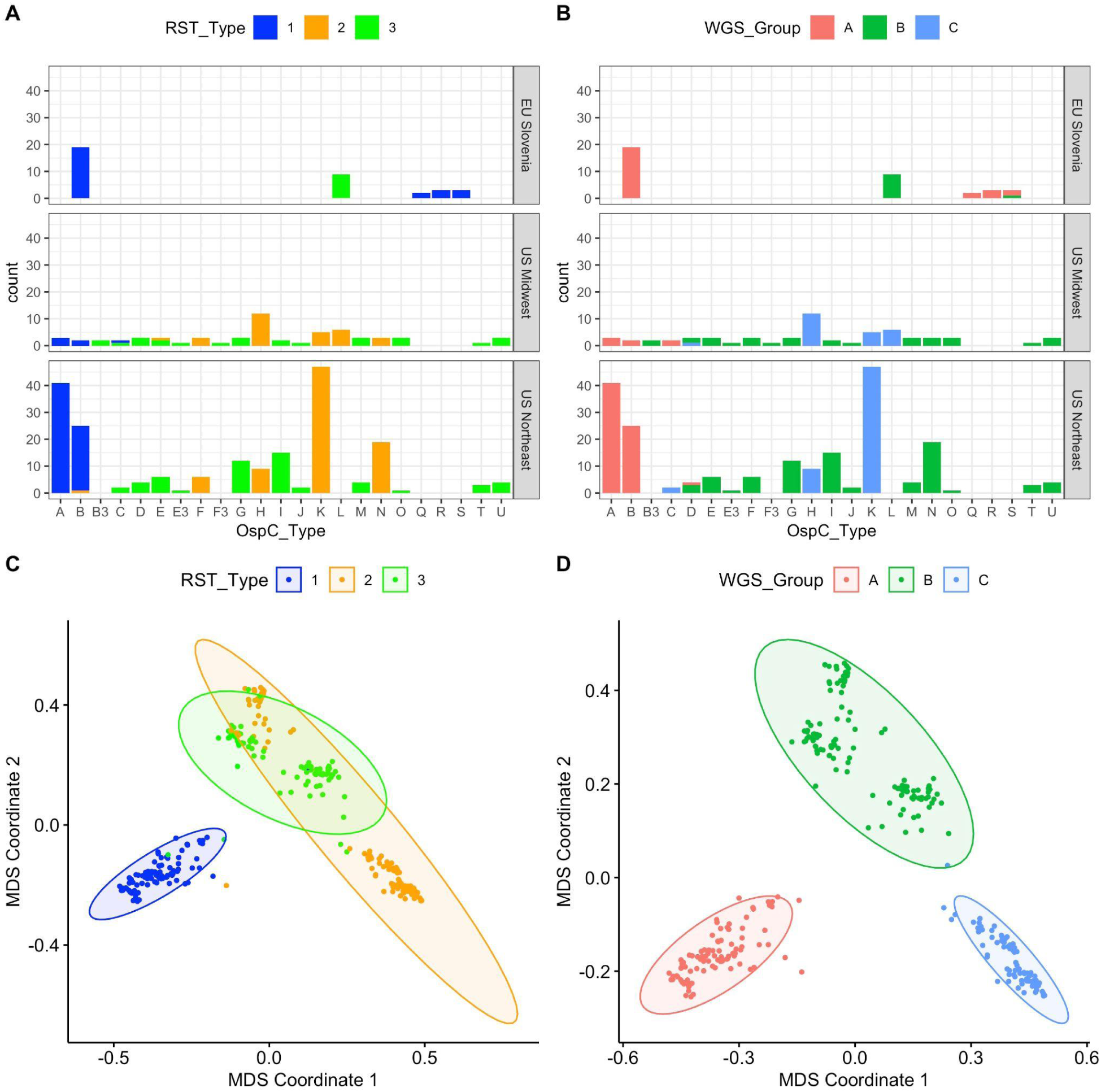
**A.** Counts of samples according to RST and OspC type. Top, middle, and lower panels show samples from different geographic regions. X-axis gives OspC type. Bars are colored according to RST type. **B.** Plots as in (A) but with bars colored according to the WGS group. **C**. Multidimensional scaling (MDS) of 299 *Bbss* genomes, with WGS RST type annotated. **D**. MDS of 299 *Bbss* genomes, with WGS type annotated.

We next constructed both maximum-likelihood (ML) and maximum clade credibility (MCC) phylogenetic trees using core genome elements (as defined by Roary[44], see methods) from WGS (Figure 2). WGS groups defined by k-mer distance corresponded to the ML clade structure on the core-genome tree and the associated OspC types (Figure 2A and E). However, they revealed substructure within these groups, particularly WGS group B, which we split into subclusters B.1 and B.2 (Figure 2B and S3B). We also inferred MCC trees using Bayesian methods as implemented in BEAST. A MCC tree is shown in Figure S2; ML and MCC trees were in broad agreement, and the posterior probability of all nodes separating WGS groups was > 0.99, indicating that the distance-based clustering was phylogenetically well-supported.

**Figure 2:**
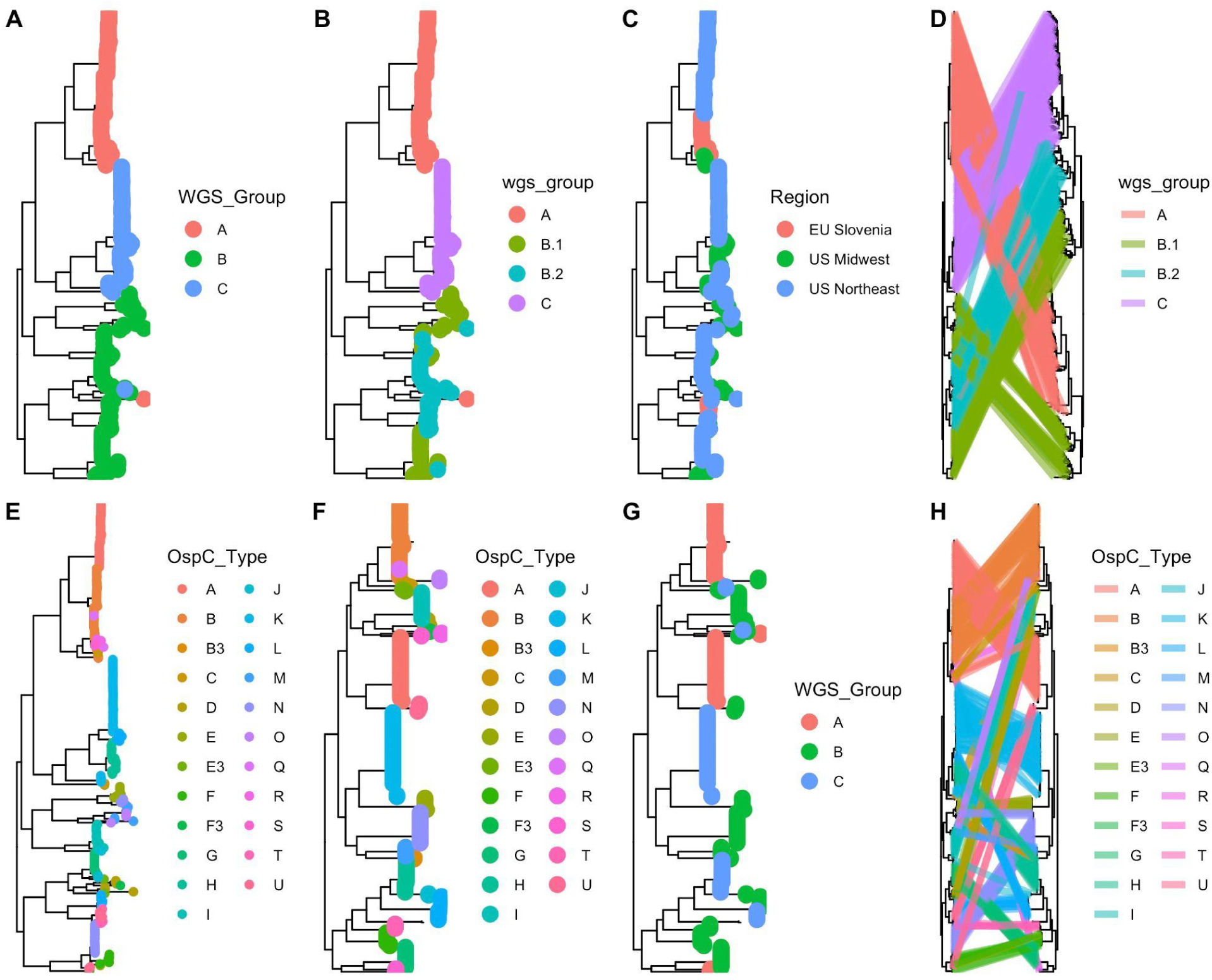
**A-B.** Core genome phylogenetic tree with tips labeled by the three major WGS groups (A). **B.** Core genome phylogeny with WGS group B split into subgroups (B.1 and B.2). **C.** Core genome phylogenetic tree with tips labeled by region of collection. **D.** The core genome phylogenetic tree (left) compared to the accessory genome phylogenetic tree (right). Lines, colored by WGS groups, connect tips from identical samples. **E.** WGS tree with tips colored by OspC type. **F.** OspC tree with tips colored by OspC type. **G.** OspC tree with tips colored by WGS group. **H.** WGS tree (left) and OspC tree with identical tips connected by strain lines, colored by OspC type.

### Comparison of Bbss isolates using classical genotyping approaches

We typed these isolates using the RST, OspC, and MLST typing schemes and compared WGS type to these existing methods (Figures 1A and 1B). Among the 299 strains, 98 were RST1 (32.7%), 112 were RST2 (37.4%), and 89 (29.8%) were RST3; 52 (17%) were OspC type K, 44 (15%) were OspC type A, 46 (15%) were OspC type B, and 21 (7%) were OspC type H. As demonstrated previously [4, 30], there was a strong linkage between RST and OspC type (Fisher’s exact test, p < 1 x 10^-6^).

In Slovenia in Europe, the most common isolates were RST1 (75%), >60% of which were OspC type B. In contrast, the most common *Bbss* isolates in the US were RST2 (41% in Northeastern US and 49% in Midwestern US), whereas RST1 strains comprised 32% of the strains in Northeastern US and only 10% in the Midwest. Further, certain OspC types have distinct geographic distributions. For example, OspC type L is found only in the Midwestern US and Slovenia and OspC types Q, R and S have only been isolated from European patients [26, 27]. These findings are consistent with previous reports that found genetic differences in *Bbss* populations based on geography [26, 27]. WGS groups were strongly associated with RST (Figures 1A-B, Fisher’s exact test, p < 1 x 10^-6^) and OspC type (Figure 1A-B, S1; Fisher’s exact test, p < 1 x 10^-6^). RST1 / Osp C type A/B sequences consistently clustered as a single clade in the core genome phylogenetic tree and MDS of k-mer distances (Figures 1C and S1), demonstrating agreement between typing methods. In contrast, RST2 and RST3 were both polyphyletic in the WGS data and contained within separate WGS groups (Figures 1C and 1D). Trees inferred from core genome sequences (Figure 2D, left panel) differed in the relatedness of major clades from those inferred from accessory genome sequences (as defined by Roary [44], see methods) (Figure 2D, right panel), but agreed on the substructure and sample membership of individual clades. This pattern, which affects major clades as a whole, indicates the occurrence of recombination events deep in the evolutionary history between core and accessory genome sequences.

Similarly, OspC types were monophyletic on the WGS tree (Figure 2E) and on a tree built from OspC sequences (Figure 2F), but WGS type was polyphyletic on the OspC tree (Figure 2G). Consistent with this polyphyly, face-to-face comparison of core genome and OspC trees demonstrated that in many cases, closely related OspC sequences were part of distinct WGS groups (Figure 2H). For example, the OspC type L isolates from the Midwestern US and Slovenia are on different branches of the core genome phylogenetic tree (Figure S2H). Thus, RST and OspC typing methods identify substructure in *Bbss* genomes, and largely agree on the divergent RST1 / OspC A/B clade. In contrast, RST does not capture fine-grain genetic structure, and OspC sequence distance does not correlate with genome-wide distance between isolates.

### Population geographic structure

We next explored the relationship between genetic markers and geography. WGS group was strongly associated with broad geographic region (US Northeast, US Midwest, EU Slovenia) (Fisher’s exact test, p < 1 x 10^-6^), similar to the findings with previously evaluated genetic markers including RST (Fisher’s exact test, p < 1 x 10^-6^) and OspC type (Fisher’s exact test, p < 1 x 10^-6^) (counts by geographic region are shown in Figures 1A-B).

Using finer-grained geographic clustering among subregions in the Northeastern US (New York, Massachusetts, Connecticut, and Rhode Island), geographic region was significantly associated with WGS group (Fisher’s exact test, p = 0.009), suggesting that geographic structuring of genotypes also occur on a regional scale (Figure S2E). The number of ORFs in the genome differed significantly by region within a given WGS group (Figure 3A). In the US Northeast and in Slovenia, WGS groups differed significantly by the number of ORFs (Figure 3B). As core genome size is relatively constant among strains regardless of geographic location, the differences in accessory genome size across different populations, even within a given genomic group with a single common ancestor, suggests that the diversification of accessory genome size may be one mechanism by which strains adapt to distinct ecological factors in each geographic region. Slovenian isolates are clustered in two well-defined monophyletic groups (Figure 2C), suggesting at least two inter-continental exchanges (Figure S2C), consistent with a previous report [15]. There were numerous (>10) exchanges between samples in the US midwest and northeast (Figure S2D).

**Figure 3:**
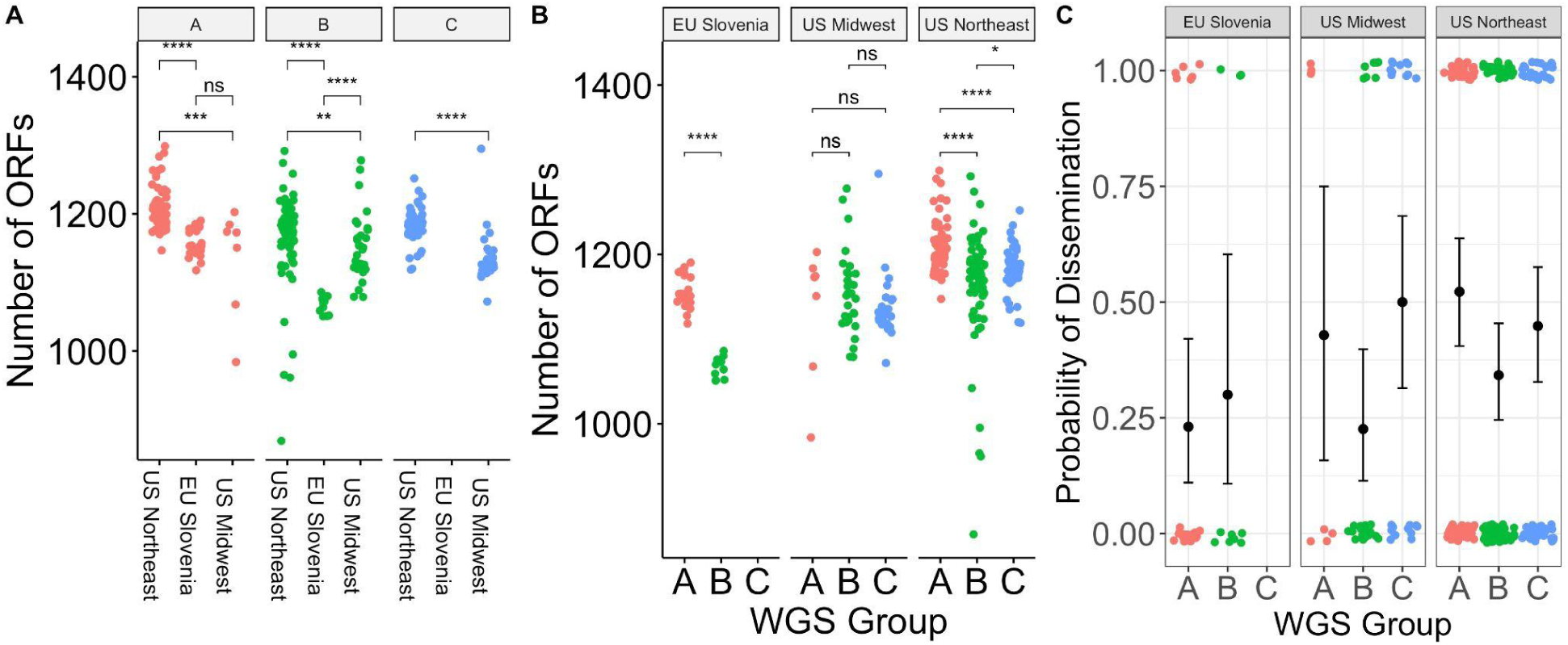
**A.** Number of ORFs by geographic region in different WGS groups. * denotes p < 0.05; ** denotes p < 0.01; *** denotes p < 0.001; **** denotes p < 0.0001; ns - not significant. **B.** Number of ORFs by WGS group in different geographic regions. **C.**Probability of dissemination by genomic group. Each point represents a sample. Points are colored by WGS group. The samples that disseminated have been plotted at y = 1; those that did not have been plotted at y = 0. Random noise has been added to the x- and y-coordinate to display the points. The mean +/- 95% binomial confidence interval is shown for each group with error bars.

We attempted to define the timing of these exchanges by inferring a time-stamped phylogeny using BEAST (Supplemental Note 1). Together, these models demonstrate a remote (hundreds of thousands to tens of millions of years) TMRCA for human-infectious strains of *Bbss*, consistent with previous estimates [52]. Precise timing requires more accurate knowledge of the mutation rate in *Bbss*.

### Associations between genotype and Bbss dissemination in patients

Dissemination is a crucial clinical event that enables the progression of disease from an EM skin lesion to more severe Lyme disease complications such as meningitis, carditis, and arthritis. Given the previously-reported associations between single-locus genetic markers and dissemination[4,5,8,30], we investigated the relationship between genotype and dissemination. We scored isolates as either disseminated or localized based on certain clinical characteristics of the patients from whom they were obtained, particularly having multiple vs 1 EM skin lesion and having neurologic Lyme disease as well as having positive culture or PCR results for *Bbss* in blood.

WGS groups differed from each other in their propensity to disseminate (p = 0.059 for 3 groups; p = 0.012 for 4 groups, Fisher’s exact test) (Figure 3C, Figure S3C). Slovenian isolates disseminated at a lower rate (25%) than US isolates (42.7%) (p = 0.045, Fisher’s exact test), and the relationship between WGS groups and dissemination was slightly stronger when testing US isolates only (p = 0.02 for 3 groups; p = 0.004 for 4 groups, Fisher’s exact test). WGS group A isolates from the US, which correlate with OspC type A and RST1 strains, showed the highest rate of dissemination (51.4%) whereas US WGS group B isolates had the lowest rate of dissemination (32.4%). Within WGS group B, there was evidence of substructure (Figure S3). US B.1 isolates disseminated at a higher rate (40.0%) than B.2 isolates (18.4%) (Figure S3C).

Consistent with previous observations [3, 4] and with the general alignment of WGS, RST, and OspC type, RST type was also associated with dissemination (p = 0.010, Fisher’s exact test), with RST1 having the greatest propensity to disseminate and RST3 the lowest [4, 5] (Figure S4B). OspC type A was also associated with dissemination (p = 0.008, Fisher’s exact test, Figure S4A). A significant association with dissemination could not be detected when OspC type was tested as a categorical variable with 23 categories (p = 0.3, Fisher’s exact test, Figure S4), but power is reduced by many categories.

The propensity to disseminate varied greatly among the US and Slovenian isolates, which is likely due to the major genetic differences in isolates between the two regions (Figure 3C). In Slovenia, the predominant WGS group A isolates are OspC type B and all the WGS-B.2 isolates are ospC type L (Figure S4). This correlation was particularly notable for WGSA strains, which were recovered from patients with disseminated Lyme disease at a rate of 51.4% in the US vs 23.1% in Slovenia. WGS-B.2 isolates in the US possess the lowest dissemination rate (18.4%), whereas those from Slovenia showed a higher dissemination rate of 30% (Figure 3D and S4A). Taken together, these data confirm that rates of dissemination vary by genotype and demonstrate that WGS A/RST1, particularly a subset distinguished by OspC type A strains, is a genetically distinct lineage with higher rates of dissemination.

### Plasmid associations with WGS profiles

As most of the genetic variation in *Bbss* occurs on plasmids [51,53,54], we investigated the variation in plasmid content across genotypes. Assembly and analysis of plasmid sequences is challenging because the length of repeated sequences in plasmids is greater than the read length generated by the short-read Illumina sequencing technology used in this study [13]. To circumvent this, we exploited the relationship between plasmid partition genes (plasmid family 32; PFam32) and plasmid types [12, 51], putatively identifying the presence or absence of a plasmid by the presence/absence of unique PFam32 sequences (Figure 4). After annotating all PFam32 genes in the assemblies using an HMM, we linked each putative PFam32 to a plasmid by finding the closest match by sequence homology from a curated list of PFam32 protein sequences (see methods).

**Figure 4:**
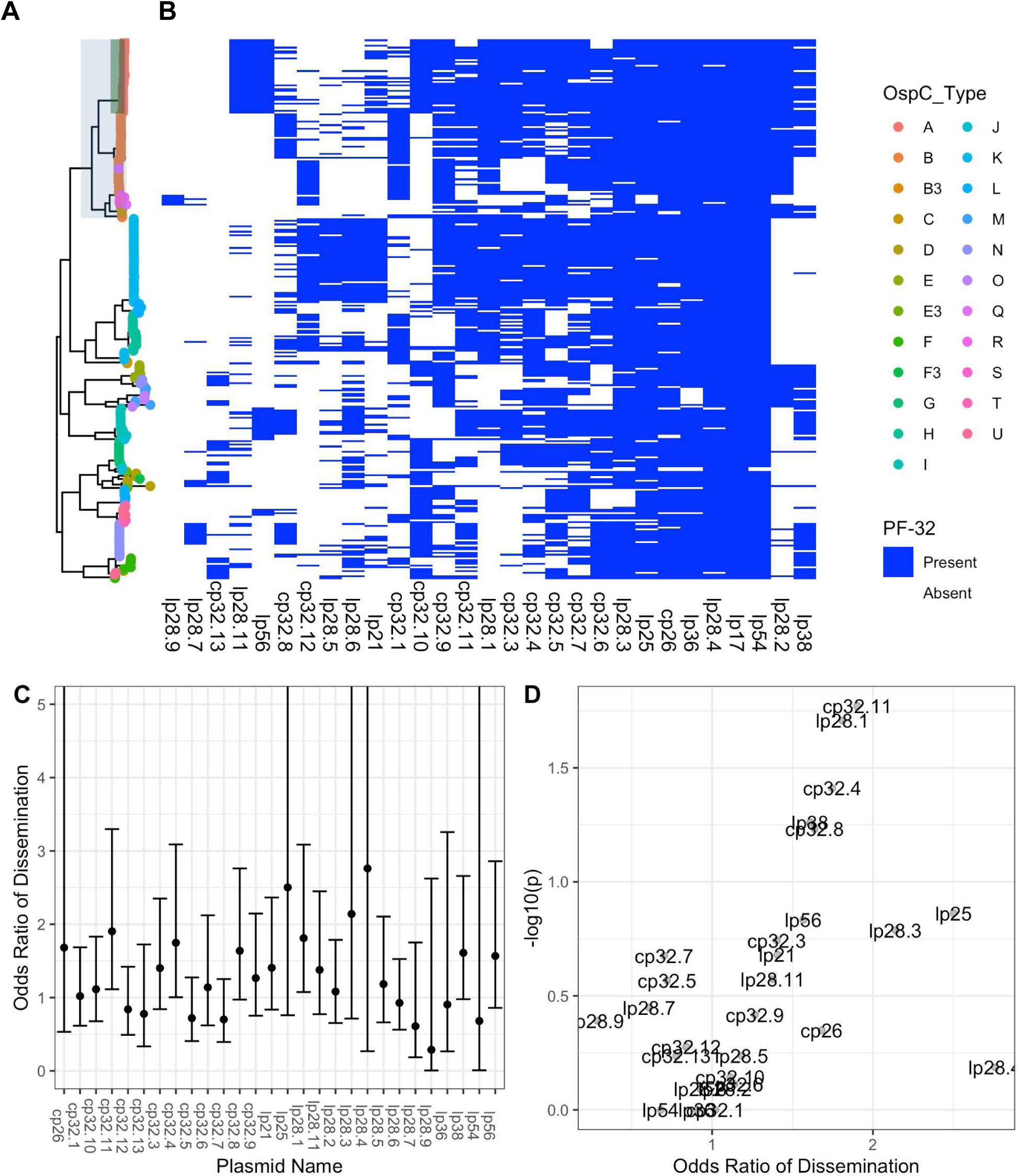
**A.** Core genome maximum likelihood phylogeny with tips colored by OspC type. The clade corresponding to RST1 is shaded in light blue and the clade corresponding to OspC type A is shaded in green. **B.** The presence/absence matrix at the right shows the presence or absence of individual plasmids using the presence or absence of Pfam32 plasmid-compatibility genes as a proxy. The columns of the matrix have been clustered using hierarchical clustering. The rows of the matrix are ordered according to the midpoint rooted maximum likelihood phylogeny shown at left. **C.** Odds ratio of dissemination and confidence interval by plasmid, inferred by Pfam32 sequences. **D.** Volcano plot displaying the -log10 P value (as calculated using Fisher’s exact test) and the odds ratio of dissemination for each plasmid, inferred by Pfam32 sequences.

Applying this method to each strain, we created a comprehensive map of plasmids across *Bbss* strains (Figure 4A-B). While a few plasmids are found more broadly, distinct genotypes and WGS groups contain unique constellations of plasmids. Several plasmids, including cp26, lp54, lp36, lp25, lp28-4, lp28-3 are found in nearly all isolates (Figure 4A-B) and others such as cp32-7, cp32-5, cp32-6, cp32-9, and cp32-3 are found in most strains. Other plasmids were more variable and only found in certain genotypes. OspC type A strains possessed a distinct plasmid profile, containing lp56 and a unique version of lp28-1 (marked by the lp28-1 PFam32 as well as a previously-annotated “orphan” PFam32 sequence, BB_F13. When found in isolation, BB_F13 defines an lp28-11 plasmid [51], so is annotated as such, although in many cases it may signify a subtype of lp28-1 rather than an entirely new plasmid (especially OspC type A isolates whose reference is likely similar to the B31 reference[9, 10]). Based on PFam32 sequences, WGS A strains also contained lp28-2 and most also contained lp38. OspC type K strains also contained a relatively homogenous subset of plasmids including lp21, lp28-5, lp28-6, cp32-12. WGS-A/ RST1 genotypes were the least heterogeneous with respect to plasmid diversity and OspC type, whereas WGS-B and WGS-C groups (RST2 and RST3) were more diverse, although the subset of RST2 strains consisting of OspC type K isolates was also relatively homogenous. Curiously, lp28-9 was found only in Slovenian RST1 isolates (Figure 4), the majority of which were OspC type B (Figure 1); cp32-12, cp32-9, and cp32-1 were also found more commonly in Slovenian isolates.

Many plasmids (e.g. lp28-1, lp28-2, lp38 and numerous others) were found in multiple distinct branches of the phylogenetic tree suggesting a complex inheritance pattern of polyphyletic loss and/or recombination. This is consistent with the previously observed reassortment between core genome elements and accessory genome elements (Figure 2D) and genetic markers such as OspC (Figure 2H). For example, OspC types B and N both contained lp28-8, whereas OspC type K genotype is most closely correlated with the lp21, lp28-5 and cp32-12 pattern. lp56 is associated with OspC type A and OspC type I.

Specific plasmids showed significant associations with dissemination. The presence of lp28-1 was associated with dissemination (OR 1.9, p = 0.01, Fisher’s exact test), as was cp32-11 (OR 1.9, p = 0.01) and cp32-4 (OR 2.0, p = 0.01) (Figure 4C-D, Supplemental Table 3). The lp38 plasmid is present in roughly half of US isolates but absent in all Slovenian isolates and demonstrated a trend toward being associated with dissemination (OR 1.6, p = 0.05).

To confirm the accuracy of these plasmid differences across genotype, we also constructed a map of plasmid occupancy across strains by an alternate approach. We aligned contigs from assembled genomes to the B31 reference sequence and annotated a plasmid as “present” if the assembled contigs covered a majority of the reference plasmid sequence (Figure S5A-C). Only plasmids present in the B31 reference genome were considered in this analysis. These results were qualitatively similar to those obtained using the PFam32 sequences (Figure S5, Supplemental Table 4) confirming that cp26, lp54, lp17, lp28-3, lp28-4 and lp36 were present in nearly all strains whereas other plasmids were more variable.

Together, these analyses reveal a core set of plasmids present across *Bbss* strains as well as strain-variable plasmids that are associated with distinct geographic and clinical features (i.e., propensity to disseminate) of *Bbss*, suggesting that they contain individual genetic elements that may underlie distinct disease phenotypes.

### Strain variation in core, accessory, and surface lipoproteome

In an effort to implicate individual genetic elements in dissemination, the core and accessory genome elements were identified in each of the sequenced isolates and all ORFs in the *de novo* assemblies were annotated and clustered using BLAST, splitting clusters whose BLAST homology was < 80% (Figure 5). Plotting the presence or absence of a given core or accessory genome element adjacent to each isolate in the phylogeny reveals consistent patterns of ORF presence/absence across closely related groups of isolates. Each of the genomic groups contained unique clusters of ORFs in the accessory genome (Figure 5). The accessory genome phylogenetic tree (Figure 2D, right) provided an alternative and more natural clustering of accessory genome elements and PFam32 sequences (Figure S6A-B).

**Figure 5:**
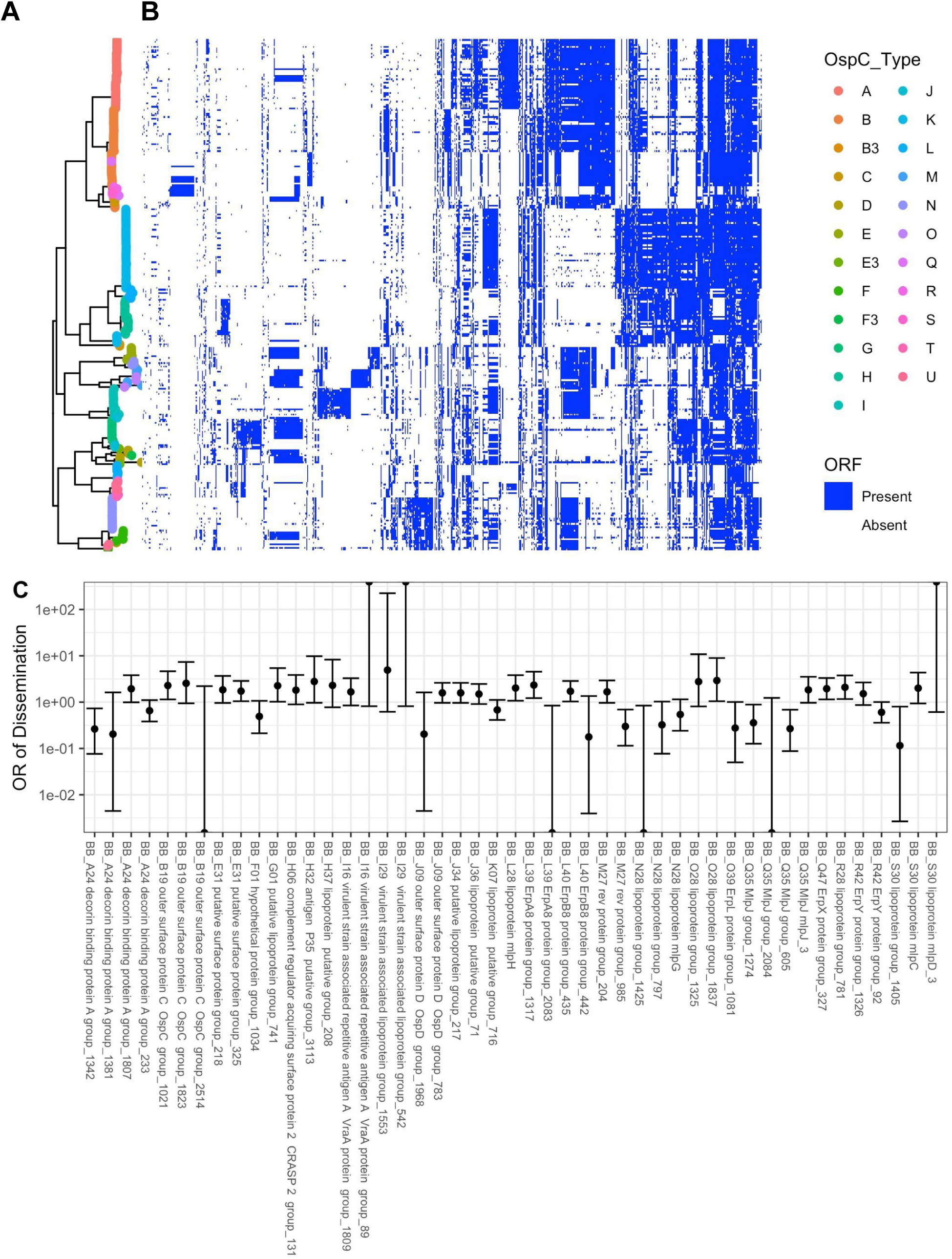
**A.** Core genome phylogeny with tips colored by OspC type. **B.** The phylogeny is plotted alongside a matrix of presence (blue) or absence (white) for genes in the accessory genome. The rows of the matrix are ordered by the phylogenetic tree in **A**. The columns of the matrix are ordered using hierarchical clustering such that genes with similar patterns of presence/absence across the sequenced isolates are grouped close together. **C.** Odds ratio (OR) of dissemination and 95% confidence interval for ortholog groups encoding surface-exposed lipoproteins and for which the unadjusted p-value for association with dissemination (by Fisher’s exact test) is < 0.15.

The most invasive genotype (WGS A) was associated with the largest pan-genome, whereas the less invasive groups (WGS Group B and C) were associated with smaller genomes (Figure 3A,B). Although many genes do not have a known function, we prioritized surface-expressed lipoproteins (Figure 6) for further analysis because of their important roles in Lyme disease pathogenesis and immunity (reviewed in [1, 55]). We focused on the subset of all lipoprotein ORFs demonstrated to be located on the surface of the spirochete [56] and divided them into core (Figure 6A) and strain-variable (Figure 6B). The *Bbss* core lipoproteome (Figure 6A) consists of approximately 45 surface lipoprotein groups that are present in almost every isolate. These include OspA and B, complement regulator acquiring surface proteins (CRASPS), as well as several other lipoproteins whose functions are less well-understood. The accessory lipoproteome (Figure 6B) consists of approximately 100 lipoprotein groups that are strain-variable. These include lipoproteins found in only subsets of isolates, such as BB_A69 and BB_E31, and others, such as Decorin binding protein A (BB_A24) and OspC (BB_B19), which were found in almost every isolate but broken into separate ortholog groups because of extensive allelic diversity. Strain-specific clusters were also present in major gene families of Erps[57, 58] (Figure S7A) and Mlps[59, 60] (Figure S7B). Larger numbers of these multi-gene family members were found in more invasive WGS groups (A and C) (Figure 6C). The number of lipoproteins in a given isolate was associated with the probability of dissemination (β_1_ = 0.037 +/- 0.017, p = 0.03, logistic regression, Figure 7D). A stronger effect was seen for Erps (β _1_ = 0.087 +/- 0.053, logistic regression, Figure 7D) with a trend toward significance (p = 0.1). In contrast, the total number of ORFs and the number of Mlp alleles were not significant in logistic regression models (p = 0.45 and p=0.38, respectively, Figure 7D). Aggregating mean effects by OspC types (Figure S7E) showed similar trends.

**Figure 6.**
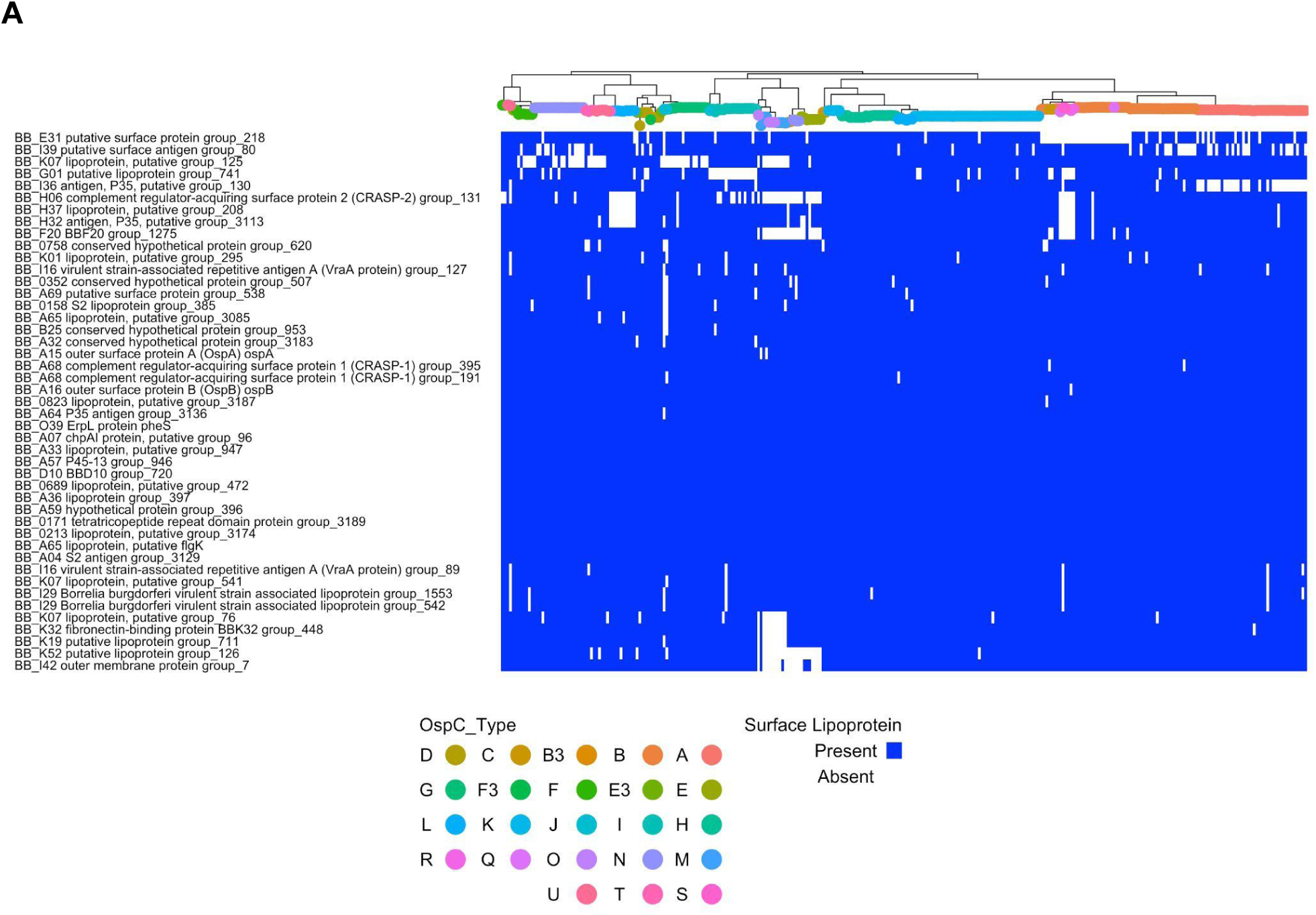
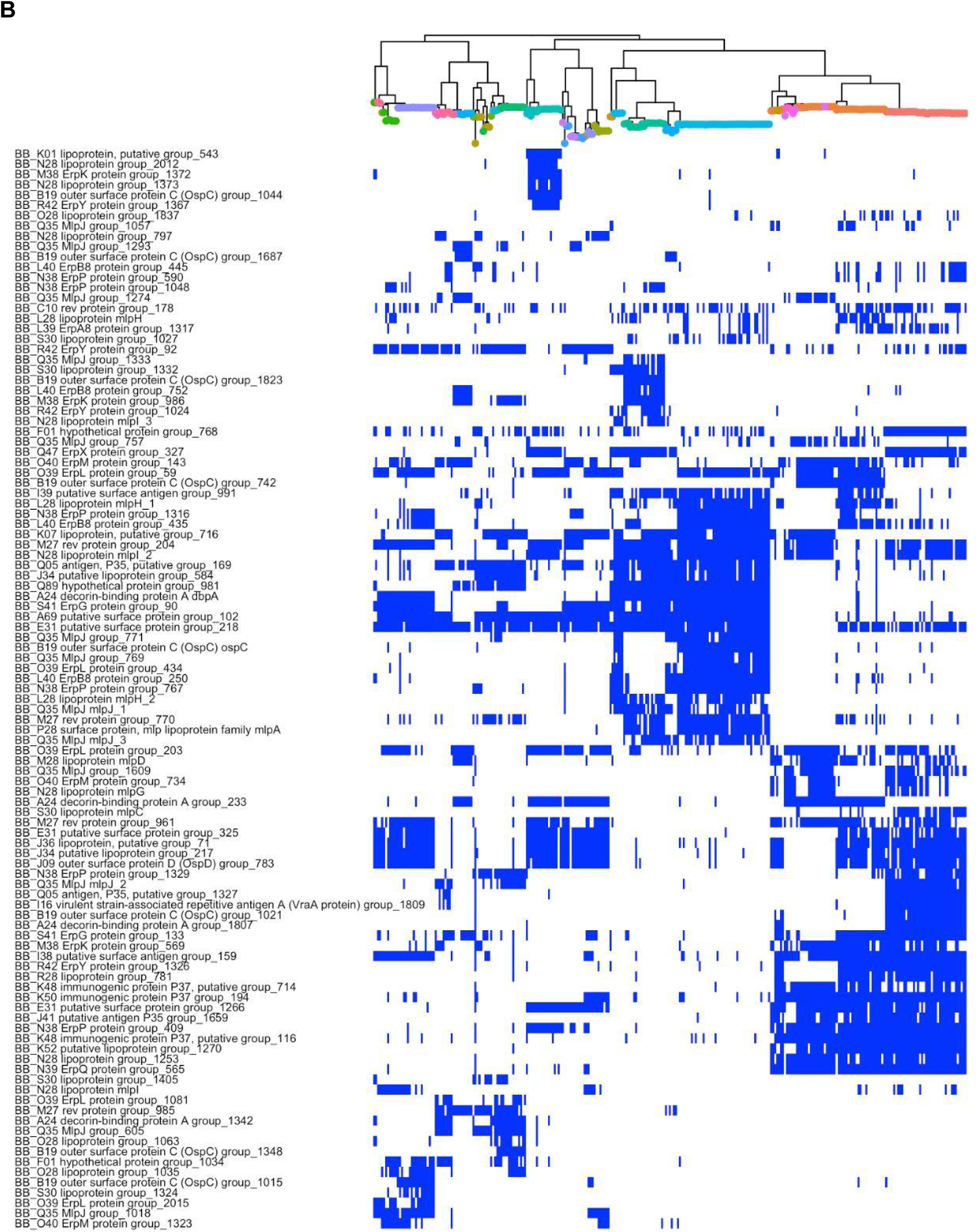
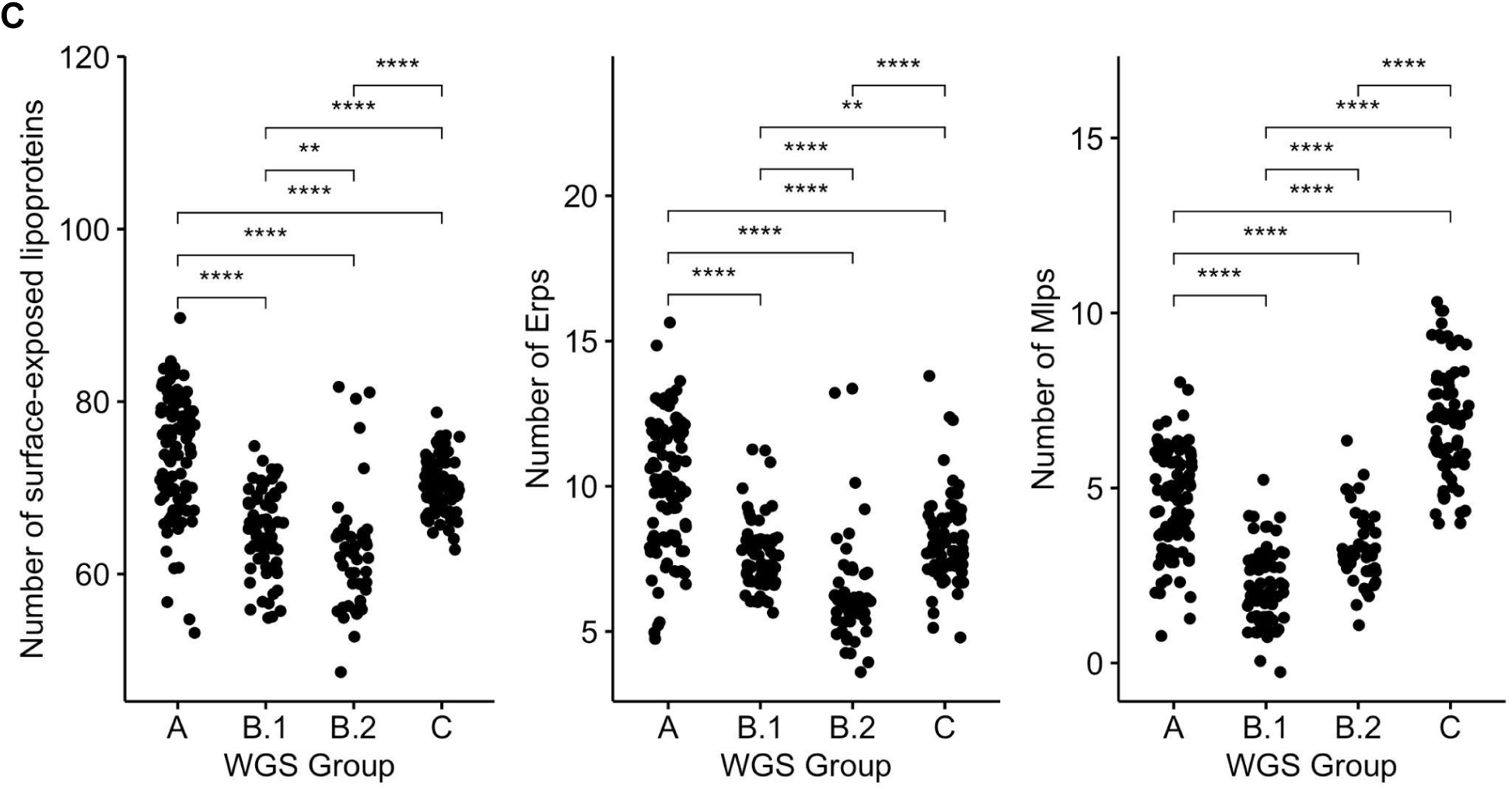
**A.** *Bbss* core surface lipoproteome: Core genome phylogeny with tips colored by OspC type (colored according to the scheme in Figure 5) with a matrix of presence (blue) or absence (white) for surface lipoproteins. Surface-exposed lipoproteins present in at least 80% of strains were considered to be part of the core lipoproteome. **B.** *Bbss* strain-variable (accessory) surface lipoproteome: Core genome phylogeny with tips colored by OspC type (colored according to the scheme in Figure 5) with a matrix of presence (blue) or absence (white) for surface lipoproteins. Surface-exposed lipoproteins present in between 5% and 80% of strains were considered to be part of the strain-variable (accessory) lipoproteome. **C.** The number of surface-exposed lipoproteins (left panel), Erps (middle panel), and Mlps (right panel) by WGS group. ** denotes p < 0.01; *** denotes p < 0.001; **** denotes p < 0.0001.

**Figure 7:**
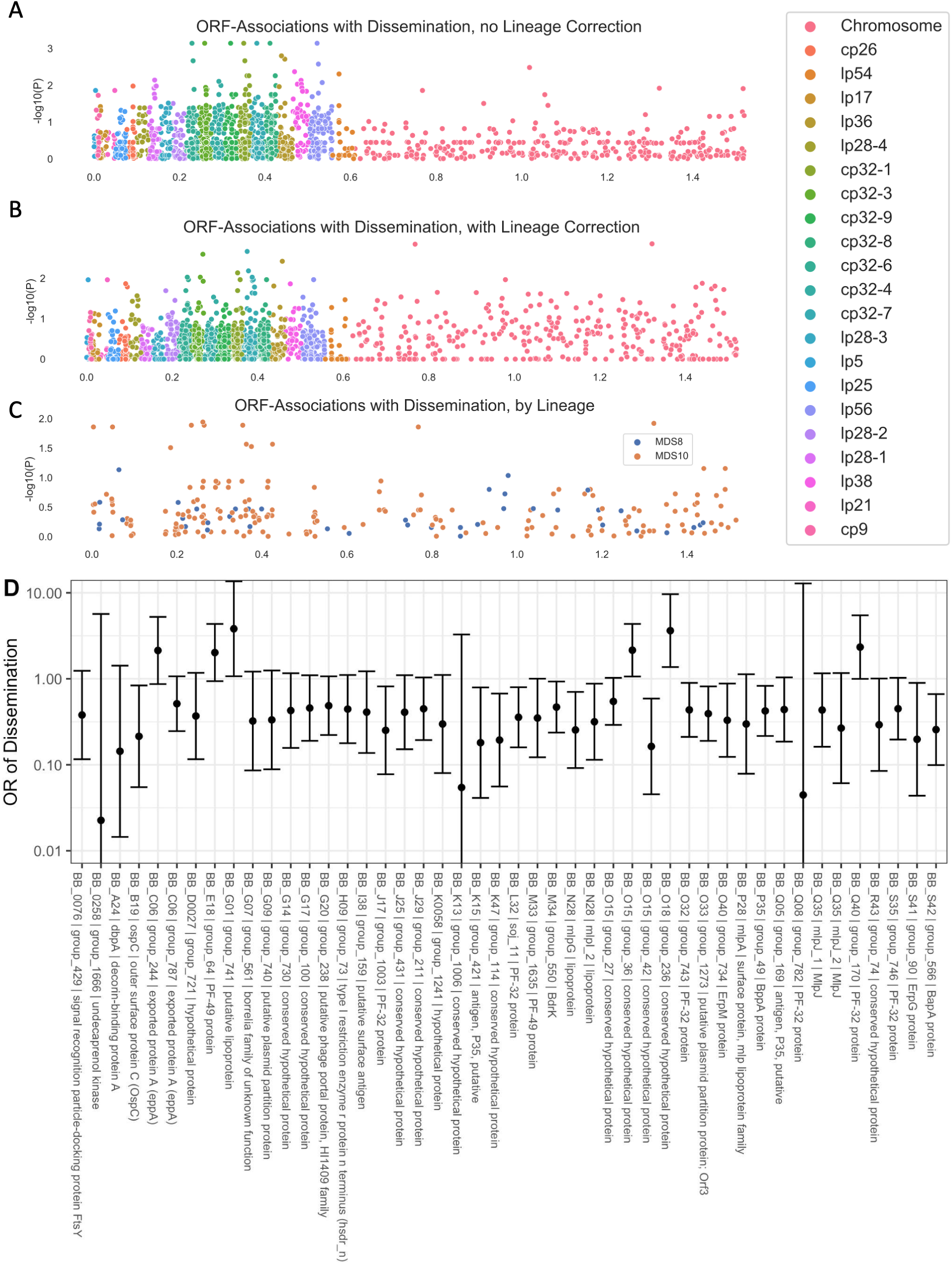
Manhattan Plots showing the association of individual ORF ortholog groups with the phenotype of dissemination. **A.** P-values from univariate logistic regression by genomic position for each ORF. **B.** P-values from regression estimates that include principal components distance matrix between strains. **C.** Manhattan plot showing loci associated with each lineage for the lineages associated with phenotype. **D.** Odds ratios (OR) (exp(beta)) with 95% confidence interval are shown for dissemination for the lineage-adjusted model. ORFs with p < 0.1 and allele frequency > 0.1 and < 0.9 are displayed.

Several lipoprotein groups, such as BBK32, BBK07, and BBK52 were found in almost all strains, but were not found in a subset of closely related genotypes. Notably, CspZ (BBH_06) and two other lipoproteins encoded on lp28-3, BB_H37 and BB_H32, were lost in two divergent subsets of Slovenian isolates (Figure 6A), suggesting multiple independent loss events in evolutionary history. Interestingly, these two subsets were either WGS-A or WGS-B.2, strains with the greatest and least probability of dissemination (Figure S3). The increased frequency of loss of lp28-3 in Slovenian isolates implies that this plasmid is likely non-essential for human infection. Moreover, this finding suggests that the selective forces acting on lp28-3 may differ in Europe and the US.

Many genes had evidence of recurrent loss or gain. For example, one cluster that shows this pattern in Figure 5B contains the lipoproteins BB_J45, BB_J34, and BB_J36 along with 12 other genes annotated on the lp38 in B31, suggesting that these lipoproteins had been lost or gained multiple times in the evolutionary tree as a part of a pattern that involved most or all of lp38.

### Associations between Accessory Genome Elements, Genotype, and Dissemination

The genetic basis of the phenotypic differences between these strains most likely includes nucleotide-level variation in chromosomal and plasmid DNA as well as variation in gene presence or absence in the accessory genome (which is primarily plasmid-borne). While it is not feasible to resolve these associations definitively in this study, we attempted to identify preliminary ORF-level associations by clustering ORFs according to homology using Roary [44]. We then applied linear mixed models genome-wide study approaches to identify ortholog groups associated with disseminated infection (Figure 7A-B). We used the approach of Earle et. al [61] to distinguish “locus” and “lineage” effects by identifying lineages that were associated with a phenotype.

Two lineages, defined by principal components of the distance matrix between isolates, were significantly associated with the phenotype of dissemination (MDS10, p = 0.02, Wald’s test), and a second component was borderline associated (MDS8, p = 0.08, Wald’s test). The results of all analyses are reported in Supplemental Table 5 and lipoprotein-specific analyses in Supplemental Table 6. In ancestry-adjusted association logistic regression analysis in which principal components were included as covariates [62], only a handful of loci were associated with phenotype, and their genomic position was distributed throughout the genome with no strong spatial pattern (Figure 7B). The uncorrected association statistics showed somewhat stronger correlations that were concentrated in the plasmids (Figure 7A).

We also used the pan-genome association approach to identify associations between ortholog groups and single-locus genetic markers. Single-locus genetic markers were strongly linked to genetic variation in ORFs, particularly among plasmids (Figure 8; Supplemental Table 7 for OspC Type A; Supplemental Table 8 for OspC Type K; Supplemental Table 9 for RST1). The strongest effects were seen among surface-exposed lipoproteins [56] (Figure S8). Together, these results, along with those of Figure 6, demonstrate that individual *Bbss* genotypes represent a tightly-linked set of genetic variation that confers a distinct surface lipoproteome.

**Figure 8:**
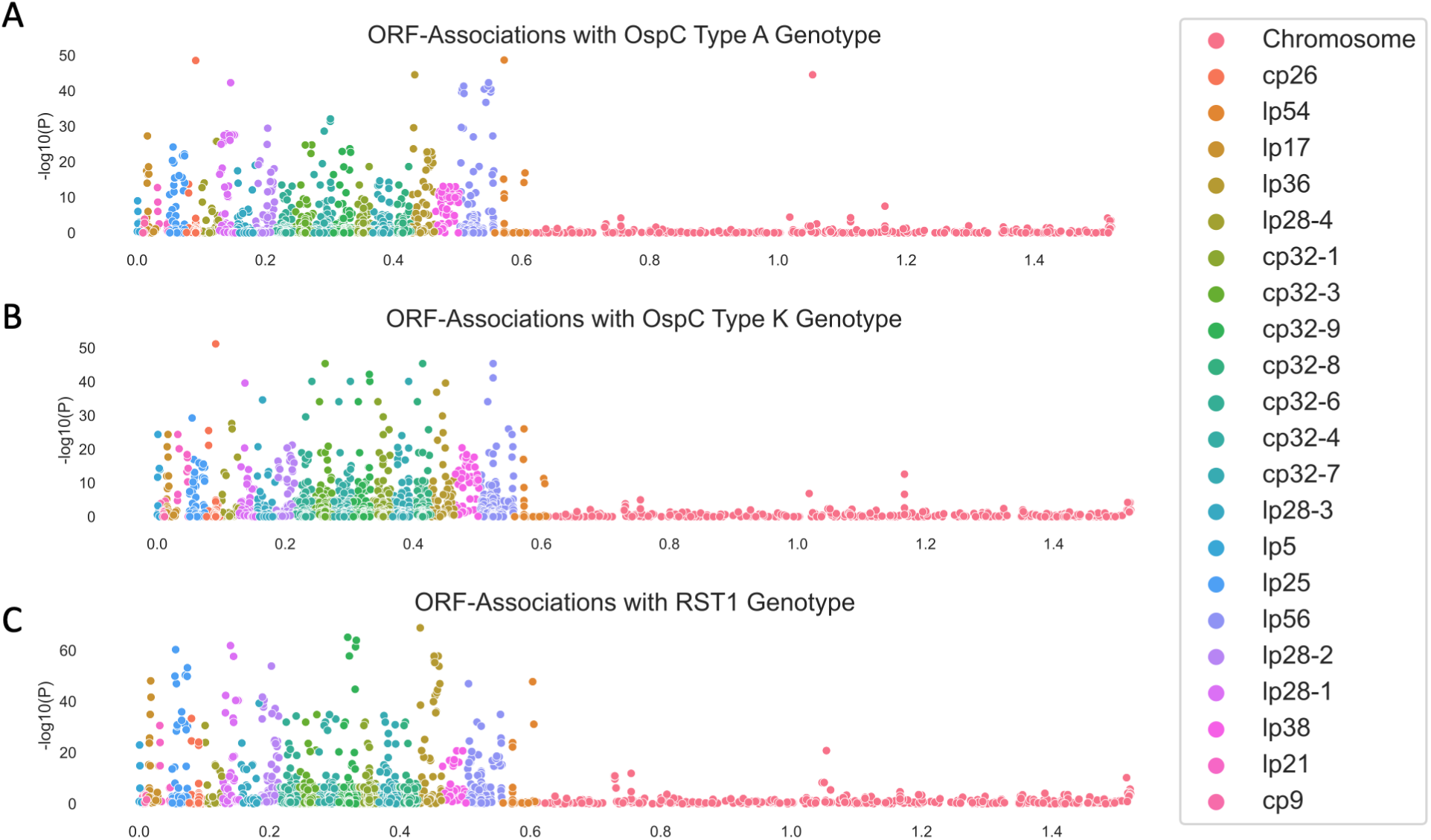
Manhattan Plots showing the association of individual ORF ortholog groups with OspC type A (panel **A**), Osp C type K (panel **B**), and RST1 (panel **C**).

Due to the structural patterns of genetic diversity in *Bbss*, ORFs associated with phenotype without ancestry correction (Figures 6D and Figure 7A) should not be ignored. Due to the near-complete linkage (e.g. Figure 8) between genetic elements in the accessory genome, individual loci with strong, causal effects on a given phenotype may not be separable from their set of linked variants, i.e. their background lineage. OspC type A strains, which are included among the strains with the highest rates of dissemination in this study (Figure S4) and as reported previously [3, 4], and which have been linked to more severe symptoms of Lyme disease [3] (Figure S4C), are strongly associated with a set of approximately 75 loci (OR > 50) including a DbpA ortholog group (OR 4964, p = 1.9 x 10^-48^, likelihood ratio test), an OspC ortholog group (OR 2951, p = 1.9 x 10^-48^, likelihood ratio test), and BB_H26 (OR 2186, p = 4.9 x 10^-38^, likelihood ratio test). These and other linked alleles were strongly correlated with one another (r = 0.94, p < 2.2 x10^-16^ for DbpA/group1807 and OspC/group1021; r = 0.85, p < 2.2 x 10^-16^ for DbpA/group and BB_H26). In many cases this linkage is physical due to presence on the same replicon (e.g. the BB_J alleles on lp38), strongly linked allelic groups may also be present on distinct replicons (e.g. DbpA on lp54 and OspC on cp26). While the strong correlations between individual alleles make it difficult to separate the statistical effects of individual alleles, such correlations are also the characteristic and defining feature of *Bbss* lineages.

## Discussion

The sequencing and analysis of 299 human clinical isolates of *Bbss* that we report here provides a previously unavailable level of resolution into the *Bbss* genetic and geographic diversity of *Bbss* strains causing Lyme disease. Our collection of WGS assemblies from these isolates—which were collected across distinct geographic regions, and which were linked to certain clinical manifestations, and systematically typed with RST, OspC, and MLST—lays a foundation for further research and advances our understanding of Lyme disease in several ways.

First, our results confirm and extend previous findings on the microbial genetic basis of disease manifestations in humans. Prior studies have identified genetic markers and correlated their presence with specific clinical findings [1,3,5,6,8,26,27,29–32], but the relationships among these markers and specific *Bbss* genes that cause phenotypic differences had not yet been studied due to limitations of existing typing systems and a lack of human isolates. Along with the novel genetic diversity uncovered by sequencing additional clinical isolates, the statistical evidence linking genetic elements to dissemination and geography that was observed in this study will be useful in prioritizing candidate genes and/or loci for further experimental evaluation. For example, we confirm here previous findings that WGS A / RST1 — particularly the subtype defined by OspC type A — is genetically distinct [27,63–65], and we identify certain genetic alterations associated with this lineage, including having a larger number of ORFs than other lineages. These ORFs are found on a strain-specific constellation of plasmids, including lp28-1 and lp56. This is consistent with previous findings that have linked the presence of lp28-1 to infectivity in mouse models [66–69]. Importantly, these results extend previous findings which showed that RST1 OspC type A strains are associated with more severe Lyme disease [3], by identifying candidate plasmids lp28-1 and lp56 as potential genetic factors associated with greater virulence of these *Bb* genotypes in patients. We show that this association, derived from mouse models, extends to humans.

Second, the microbial genetic association studies presented here begin to resolve the individual genetic elements underlying certain human phenotypes of Lyme disease. Using two different methods to infer the presence or absence of plasmids, we provide the first plasmid presence / absence maps of a large collection of human clinical isolates. Integrating this information with associations at the level of individual ORFs provides a clearer view of the potential determinants of distinct phenotypes. While we cannot yet resolve the causative loci on lp28-1 or lp56 that enhance the pathogenicity of OspC type A strains, we highlight candidate loci and quantify the statistical evidence for each locus considered. ORFs in these plasmids such as BB_Q67 (which encodes a restriction enzyme modification system [70, 71]), BB_Q09, BB_Q05, BB_Q06, BB_Q07, BB_J31, BB_J41 and others (Supplemental Table 8) are tightly linked to the OspC type A genotype and are candidates for further experimental study.

In addition, our sub-analysis of surface-exposed lipoprotein sequences (Figures 6A and 6B) may also be useful for experimental follow-up given the importance of surface lipoproteins for immunity, pathogenesis, and *Bbss-*host interactions (reviewed in [1, 55]). Particular alleles of DbpA (BB_A24), and specific members of the Erp (BB_M38, BB_L39) and Mlp (BB_Q35) (supplemental data file 2, Figures 6C and 6D) families are associated with dissemination and represent potential candidates for evaluation in follow-up studies. Both the specific list of ORFs strongly associated with OspC type A and the more general pattern of variation across strains provides clues into enhanced virulence. Among those ORF groups associated with the OspC genotype, allelic variants of DpbA have been shown to promote dissemination and alter tissue tropism in a mouse model of Lyme disease [72]. Multiple genes in linked blocks probably contribute to pathogenesis. For example, In OspC type A strains, DbpA is strongly linked to OspC type A. Allelic variation in OspC alters binding to extracellular matrix components, promotes joint invasion, and modulates joint colonization[73]; OspC has also been shown to promote resistance in serum killing assays [74], and its role in causing infection can be, under certain circumstances, partially complemented by other surface lipoproteins [75, 76].

Our data suggest that copy number among multi-copy gene families may be linked to dissemination. Given that Erps are divided into three families that each bind to distinct host components (extracellular matrix, complement component, or complement regulatory protein) [58,77–80]; it is possible that the strain-variable clusters of Erps (Figure S7B, Figure S7D-E) may influence clinical manifestations by modulating strain-specific properties of tissue adhesion or resistance to complement-mediated killing of spirochetes. The functions of Mlp proteins and many other strain-variable lipoproteins are still not well understood. The statistically-significant relationship between lipoprotein number and probability of dissemination and the borderline-significant relationships for copy number of Erps and Mlps (Figure S7D-E) suggest that varying the amount and diversity of linked clusters of surface lipoproteins—which, individually or in combination, may promote survival in the presence of immune defenses, binding to mammalian host tissues and other pathogenic mechanisms— may be a general mechanism for strain-specific virulence of *Bbss*.

Using unadjusted, univariate associate models, virtually all dissemination-associated genes were found on plasmids. However, after correction for spirochete genetic structure due to lineage, only weak locus-specific associations were observed. The block patterns of Figures 5 and 6 demonstrate why this is the case. Genes are inherited in blocks; the inheritance pattern of genes within these blocks is strongly correlated such that only infrequently are genes from within a block found in isolates that are outside the block. This pattern is also seen in plasmids, and plasmids are a natural mechanism for this pattern of inheritance. An important consequence of this finding is that it may not be possible to resolve individual loci beyond correlated blocks of genes simply by increasing the number of samples or other methods to improve statistical power because the near-complete correlation between individual loci makes it statistically difficult to distinguish the individual effects among correlated genes. Thus, beyond identifying genomic elements or groups of correlated genes associated with a phenotype, further fine mapping will require biological experiments with reverse genetic tools. The results shown in Figure 6 and 7, and Supplemental Data File 3 are helpful in narrowing down the candidate loci and genetic elements that may predispose to or protect from dissemination.

Third, our analysis highlights how evolutionary history, geography, and differences in strain genetic diversity interact in complex ways to contribute to clinical heterogeneity in Lyme disease. In the context of known associations between genotype and clinical disease, the difference in genetic markers across geographic areas may help explain why some clinical phenotypes are more common in certain geographic locations. For example, Lyme arthritis is more common in the US compared to Europe, probably because the infection in the US is due predominantly to *Bb*ss strains which are more arthritogenic [81]). OspC type A strains appear to be more common among patients in the US Northeast [26, 27]. The intermixing of WGS groups B and C in RST types 2 and 3 has not been a major issue in practice because the phenotypes (for example, the relative rate of dissemination) of those groups appear more similar than the genomically and phenotypically divergent RST1 / WGS A group. Similarly, OspC genotyping has its limitations. The large number of OspC types (at least 30) makes phenotypic associations with specific OspC genotypes challenging. More importantly, the discordance between OspC sequences and whole-genome phylogenies—a discrepancy observed since the earliest OspC sequences were published [82] and likely related to the fact that the OspC locus is a known recombination hotspot on cp26 [83]—may make OspC unreliable as a genetic marker of phenotypic traits. In this regard, WGS serves as a gold standard against which other typing methods can be compared, facilitated here by our sequenced and fully-typed set of isolates.

WGS also offers new insight into evolutionary history and population divergence of *Bbss.* Estimation of divergence times suggests a remote (at least hundreds of thousands of years) origin for human infectious *Bbss.* The similarity in TMRCA estimates for samples from Slovenia, the US Northeast, and the US Midwest indicate that the common ancestry for sequences currently circulating in these populations is also remote; however, the strong lineage structure and history of multiple exchanges suggests that the local history of distinct lineages is also complex likely with multiple inter-region migration events. The consistent directional differences in ORF number by region also suggest that adaptive evolution to local environments has occurred, exploiting mechanisms of gene loss/gain on plasmids.

This report has several limitations. First, plasmids pose a unique challenge for assembly and annotation [10, 12]. As others have shown [13], complete plasmid assembly with short read sequences is not possible. We devised two bioinformatic methods to overcome these changes and infer plasmid presence/absence from short read sequencing, but neither is perfect. Our PFam32 analysis is limited by an uncertainty as to which gene sequences are contained on the plasmid associated with the PFam32 sequence. A complementary analysis based on the B31 reference sequence relies on a high-quality pre-existing assembly but cannot account for genes/plasmids absent from the B31 reference. We also cannot exclude the possibility of plasmid loss during culture, but isolates were passaged fewer than five times to minimize this possibility.

Second, there are limitations due to analysis of isolates collected over time by different groups at different sites. In particular, we may underestimate dissemination because an assessment of spirochetemia (blood PCR or blood culture) was only available for 70.9% of isolates (supplemental data file 3) and the absence of positive culture or blood PCR from a single time point does not rule out the possibility that dissemination from the initial skin lesion may have occurred or may occur at a later time in untreated patients.

Third, there are statistical limitations related to the *Bbss* genome and study size. Models that naively correlate a given gene with the phenotype of interest will produce spurious associations due to the confounding effect of lineage and may overstate the effect from single loci, a problem which is well known in human genome-wide association studies [84]. Corrections for lineage and population structure are often applied to human [85, 86] and bacterial [61, 62] association studies. However, *B. burgdorferi* underscores the challenges to these approaches, both because lineages appear to be *defined* by the exchange of blocks of genes and because the coarse tree structure differs for the core and accessory genomes, implying that a single similarity measure to capture the pairwise dissimilarity between strains may not be adequate. Larger studies with more isolates, statistical methods that incorporate the joint distribution between genetic markers, and plasmid assemblies finished by long read sequencing are required as a next step. The present study includes isolates collected by different investigators over the past 30 years. Due to the logistical complexity and cost of collecting *Bbss* isolates from patients in clinical studies, substantially larger studies of *Bbss* from patients may not be feasible in the near term; however, long-read sequencing approaches have improved in accuracy, availability, and cost, making finishing the genomes of existing isolates a logical next step.

Taken together, our results indicate that each *Bbss* genotype represents a tightly-linked constellation of strain-specific variation that occurs primarily in plasmids, much of it involving surface-exposed lipoproteins. OspC type A strains—with their enlarged pan-genome, distinct set of plasmids, including lp28-1 and lp56, and variants of many surface lipoproteins, particularly a unique subtype of DbpA—represent the most dramatic example of this genetic signature associated with distinct phenotypes of Lyme disease in humans. Nevertheless, the pattern is generalizable across genotypes and, given the strong linkage between microbial genotype and phenotype for *Bbsl,* and similarities in genetic structure among *Bbsl* genomes, is likely true broadly across all Lyme disease agents (*Bbsl*).

## Acknowledgments

This work was supported by a Doris Duke Charitable Foundation Physician Scientist Fellowship (2019123 to J.E.L), the National Institute of Allergy and Infectious Diseases (K99/R00148604 to J.E.L, U19AI110818 and U01 AI151812 to P.C.S.; R01AI045801 to I.S., and R21AI144916 to K.S.), the National Institute of Arthritis and Musculoskeletal and Skin Diseases (R01AR41511 to I.S. and K01AR062098 to K.S.), the Bay Area Lyme Foundation (to P.C.S. and J.E.L.), the Howard Hughes Medical Institute (P.C.S.), the Arthritis Foundation Fellowship (to K.S.), and the Slovenian Research Agency (P3-0296, J3-1744, and J3-8195 to F.S.).

## Data and code availability

Genome sequences reported here have been deposited in Genbank under PRJNA923804. Code is available at https://github.com/JacobLemieux/borreliaseq.

## Supplemental Figures

**Supplemental Figure 1:**
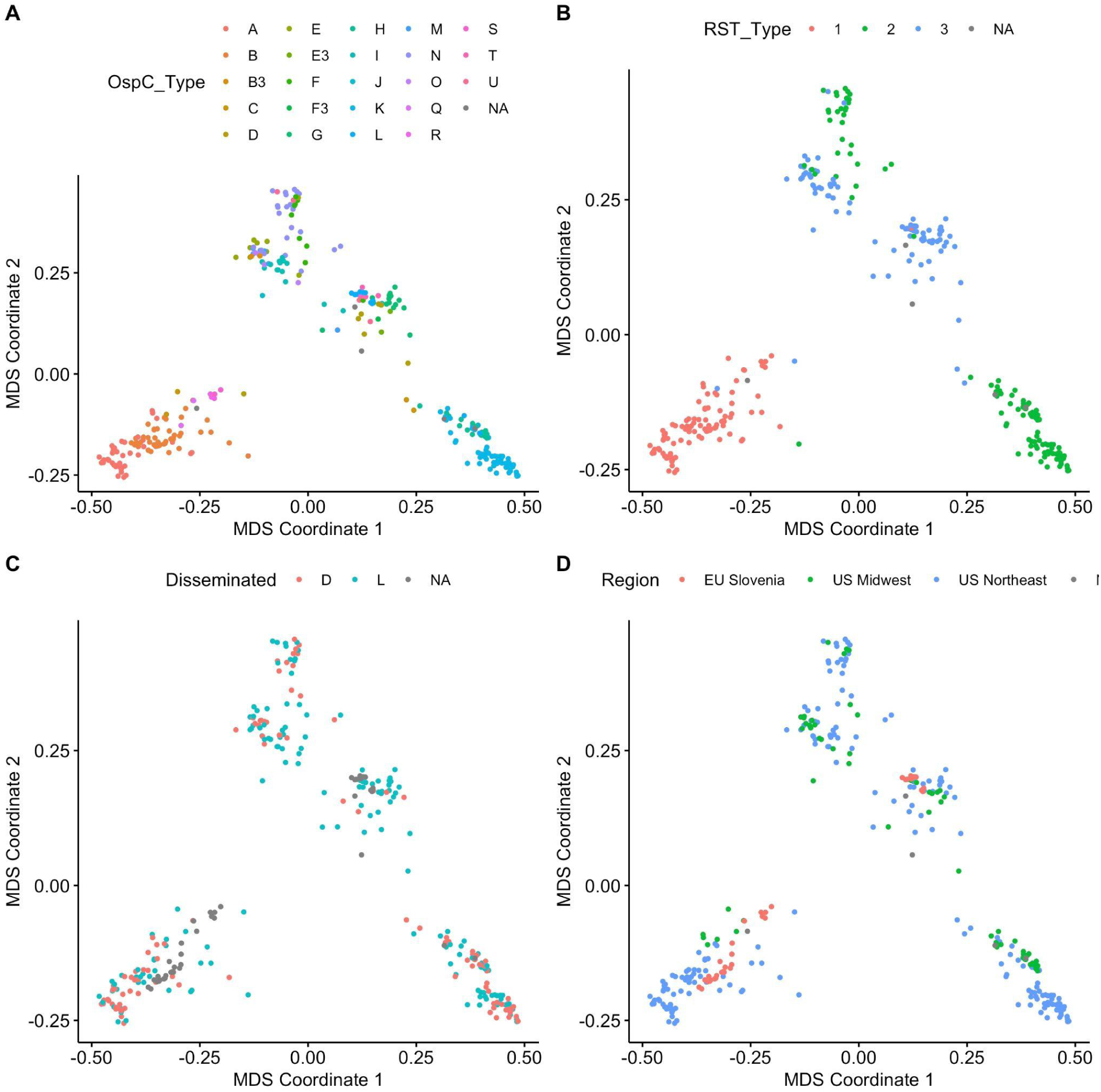
Multidimensional scaling (MDS) reveals the population structure of US and Slovenian *Bbss* isolates.

**Supplemental Figure 2:**
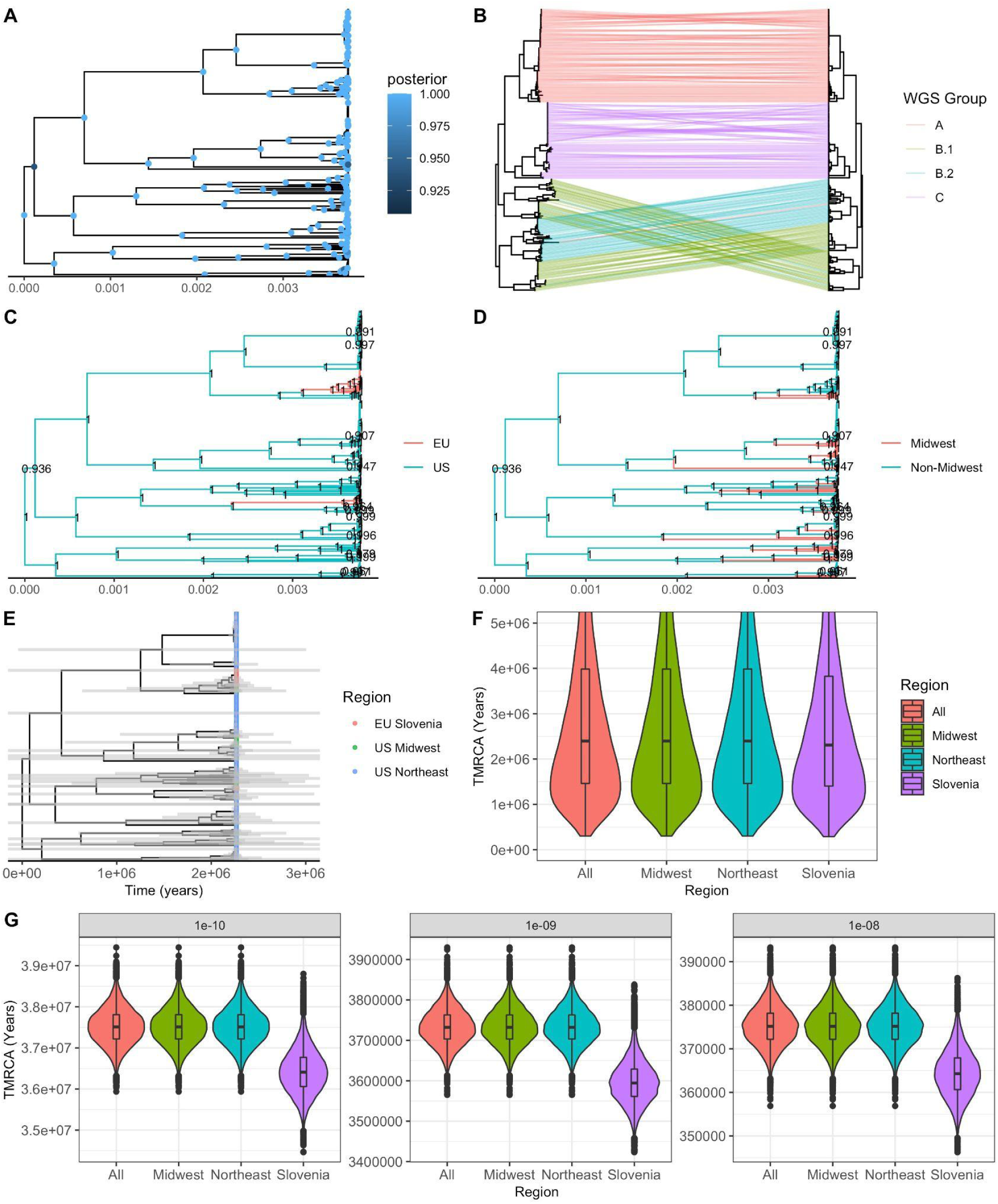
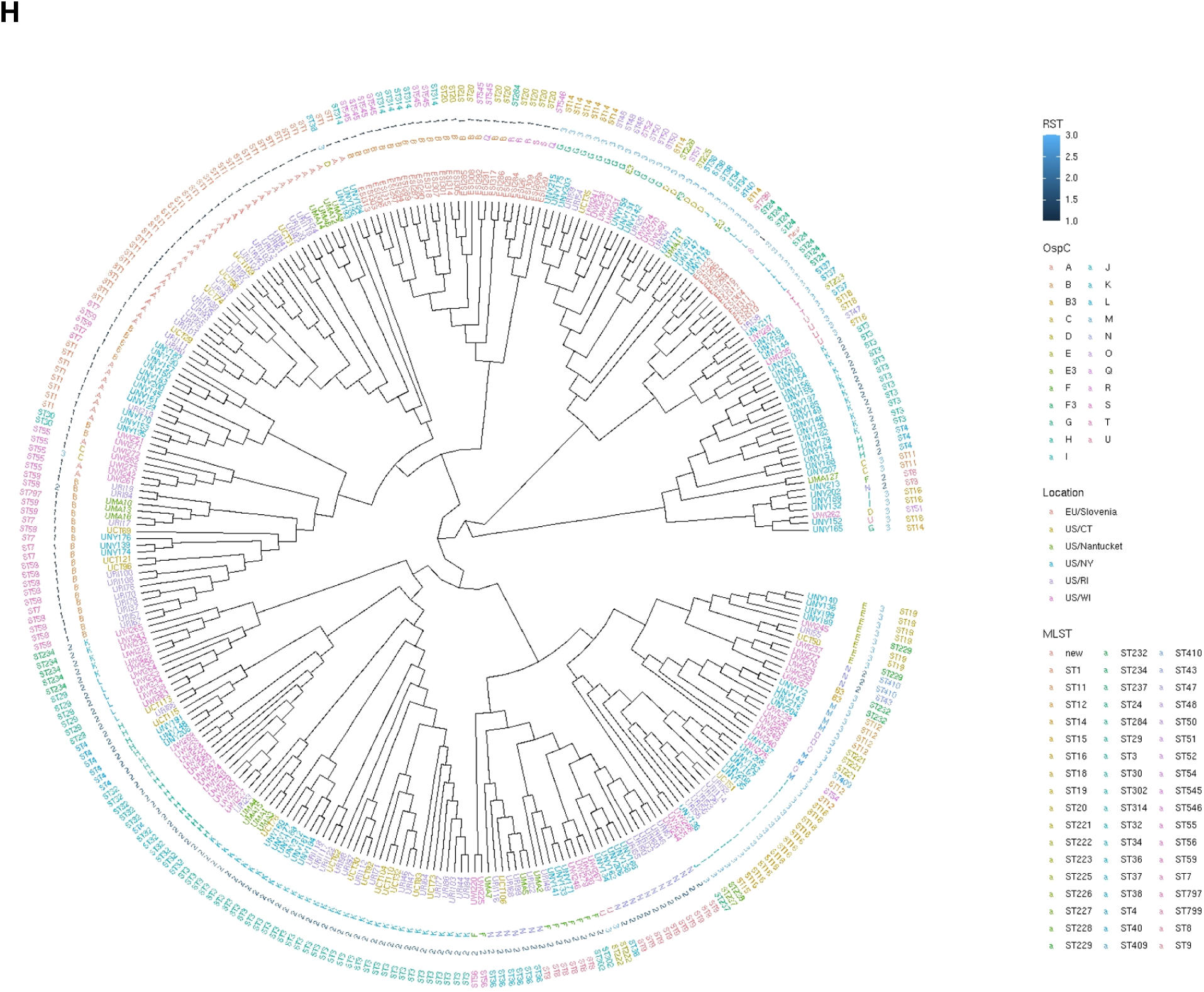
**A.** Maximum clade credibility (MCC) tree. Nodes with posterior probability > 0.9 are colored. **B.** Maximum likelihood (left panel) and MCC tree, with identical tips connected with lines colored according to WGS group. **C.** MCC tree with nodes with posterior probability > 0.9 labeled. Tips from the US have been grouped and their most recent common ancestor are colored blue; all others are colored red. **D.** MCC tree with nodes with posterior probability > 0.9 labeled. Tips from outside the US Midwest have been grouped and their most recent common ancestor are colored blue; all others are colored red. **E.** Time-tree with 95% credible interval of node heights plotted as gray bars. **F.** Density of time to most recent common ancestry (TMRCA) for major subpopulations and the full sample set (root). An inset boxplot gives the median and IQR. **G.** Density of time to most recent common ancestry (TMRCA) for major subpopulations and the full sample set (root) under three different fixed-clock models with the clock rate set at 1×10^-10^ substitutions/site/yr (left panel), 1×10^-9^ substitutions/site/yr (middle panel), or 1×10^-8^ substitutions/site/yr (right panel). **H.** Core genome phylogeny of 299 whole-genome sequences. The phylogeny is shown as a cladeogram (branch length does not correspond to genetic distance). The tips are labeled with sample names. RST type, OspC type, location, and MLST type are annotated. Whole genome sequences recapitulates existing typing schemes while adding additional resolution. Geographic origin is associated with different branches of the tree. For example, Slovenian isolates cluster in two distinct branches.

**Supplemental Figure 3:**
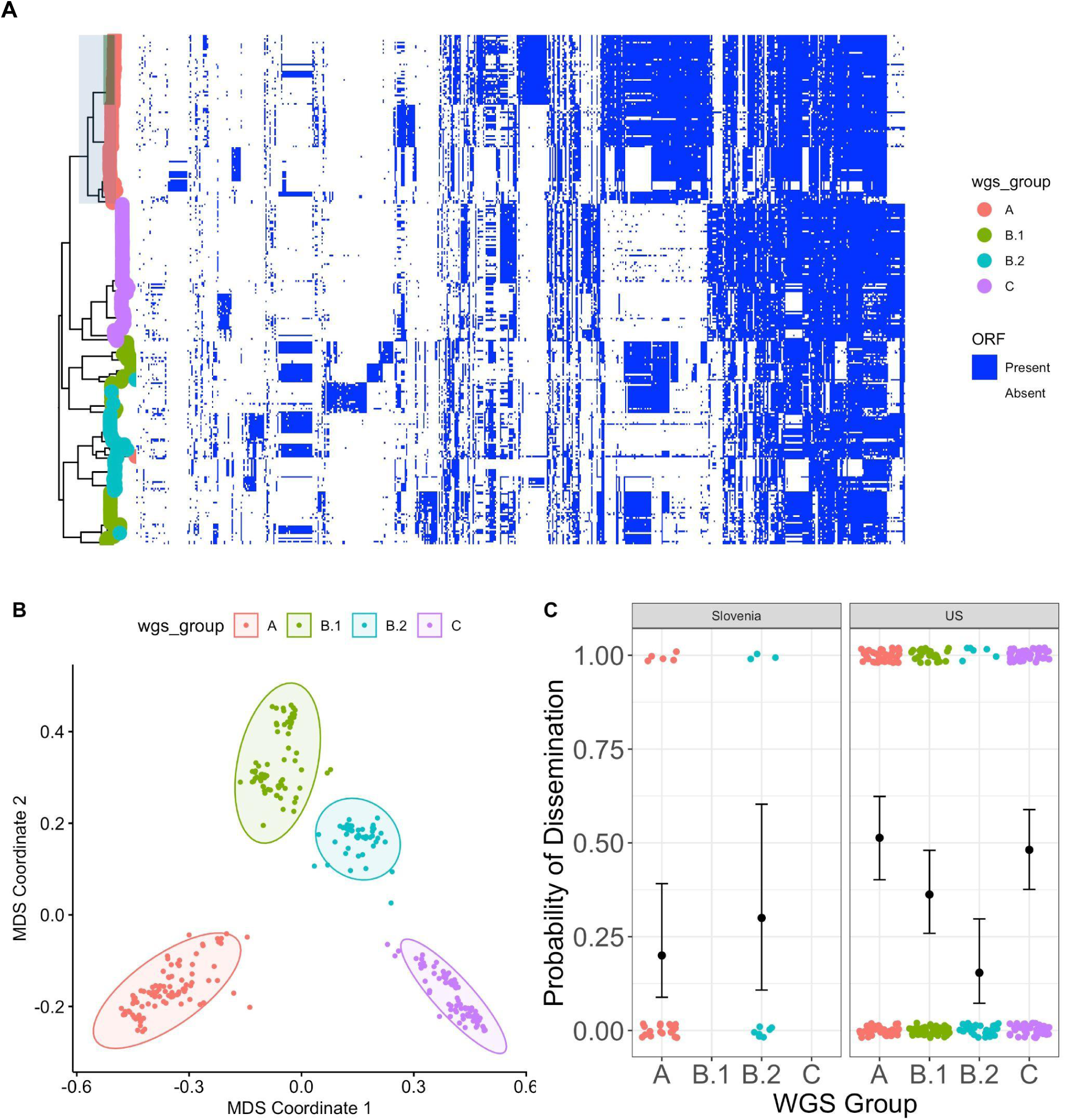
**A.** Core genome phylogenetic tree colored by WGS groups A-C with group B divided into B.1 and B2; accessory genome presence/absence matrix is reproduced from Figure 5 to highlight accessory genome elements that correlate with B.1 and B.2 sublineages. The clade corresponding to RST1 is shaded in light blue and the clade corresponding to OspC type A is shaded in green. **B.** MDS plot with group B divided into B.1 and B.2. **C.** Probability of dissemination by genomic group using the four groups including B.1 and B.2.

**Supplemental Figure 4:**
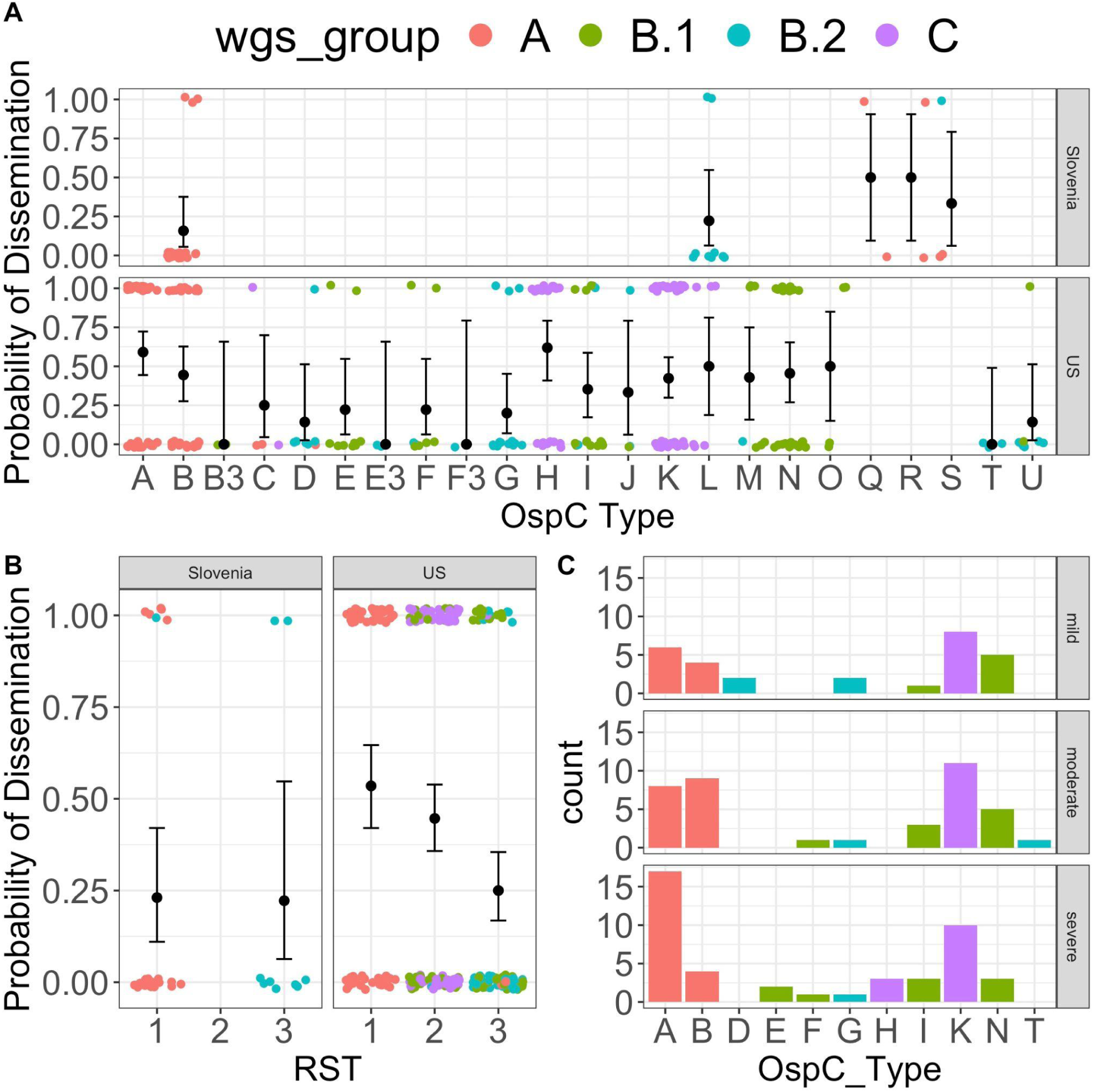
Probability of dissemination by **(A)** OspC type and **(B)** RST. **C.** Severity of Lyme disease by OspC type with WGS group shown by color.

**Supplemental Figure 5:**
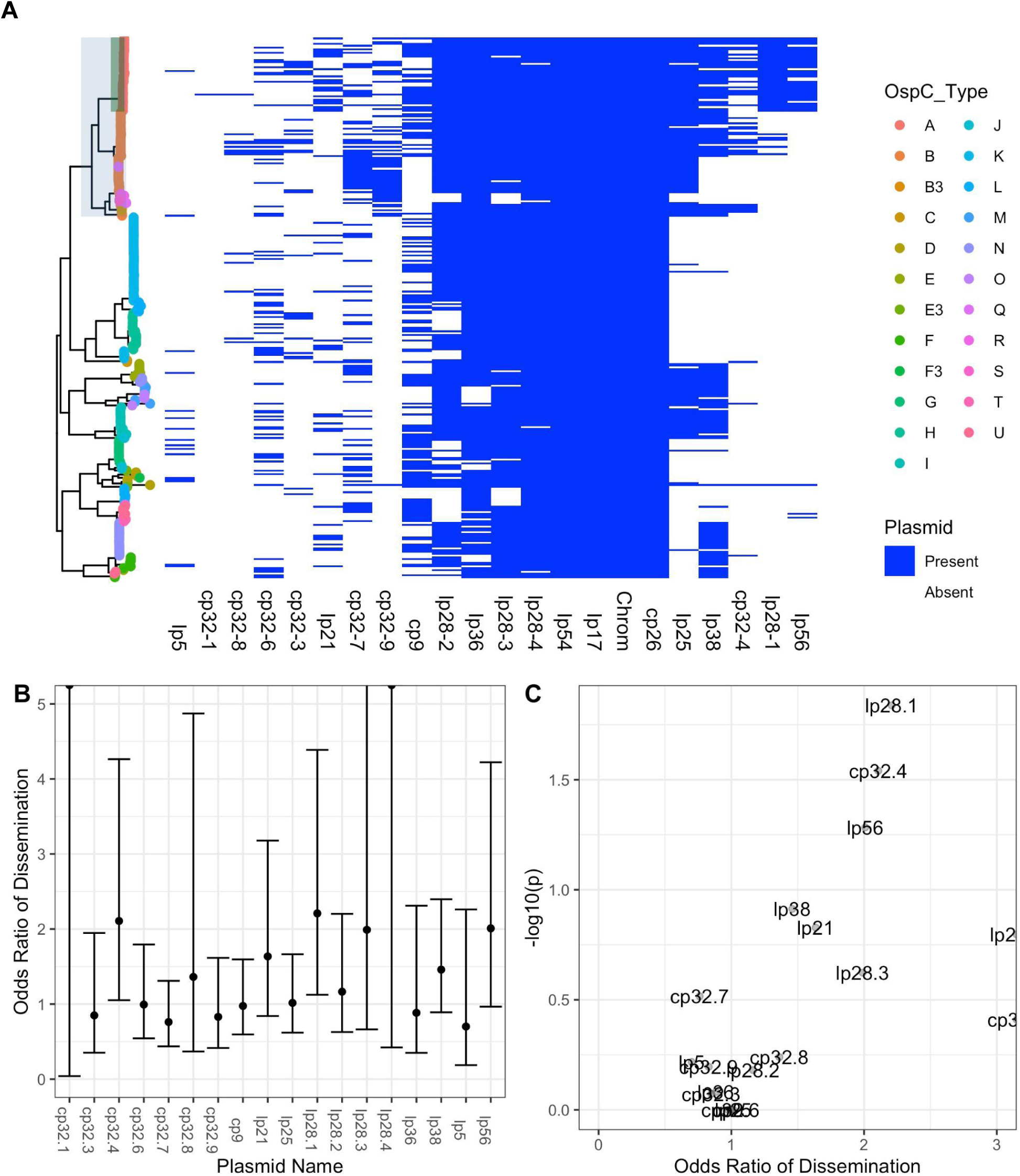
**A.** Inferred presence / absence of a plasmid based on alignment of assembly contigs to the B31 reference. A plasmid is inferred as ‘present’ in the isolate if > 50% of the length is covered by aligned contigs in the de novo assembly for the genome of the corresponding isolate. The clade corresponding to RST1 is shaded in light blue and the clade corresponding to OspC type A is shaded in green. **B.** Odds ratio of dissemination and confidence interval by plasmid, inferred by Pfam32 sequences. **C.** Volcano plot displaying the -log10 P value (as calculated using Fisher’s exact test) and the odds ratio of dissemination for each plasmid, inferred by alignment of assembled contigs to the B31 reference sequence.

**Supplemental Figure 6:**
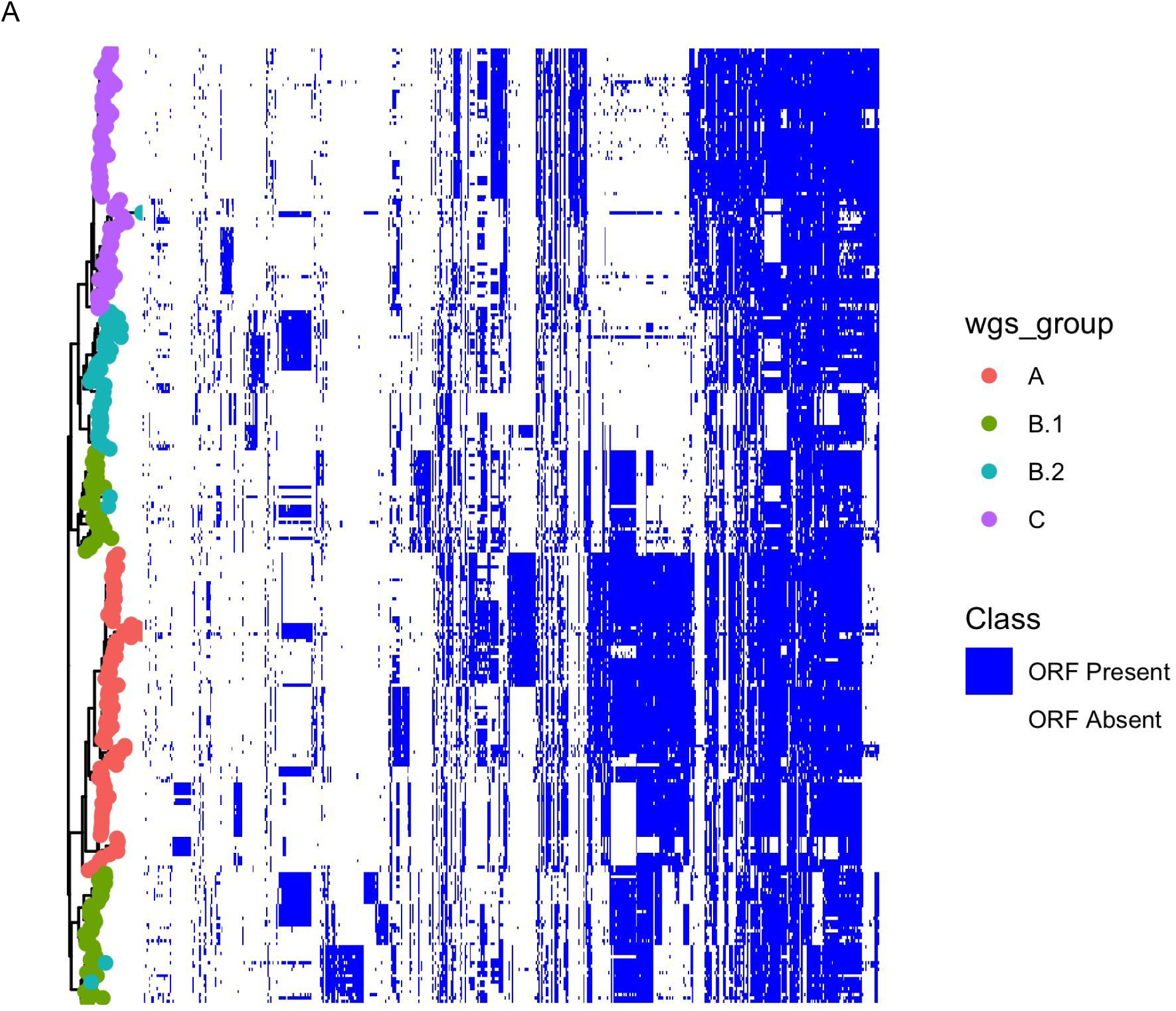
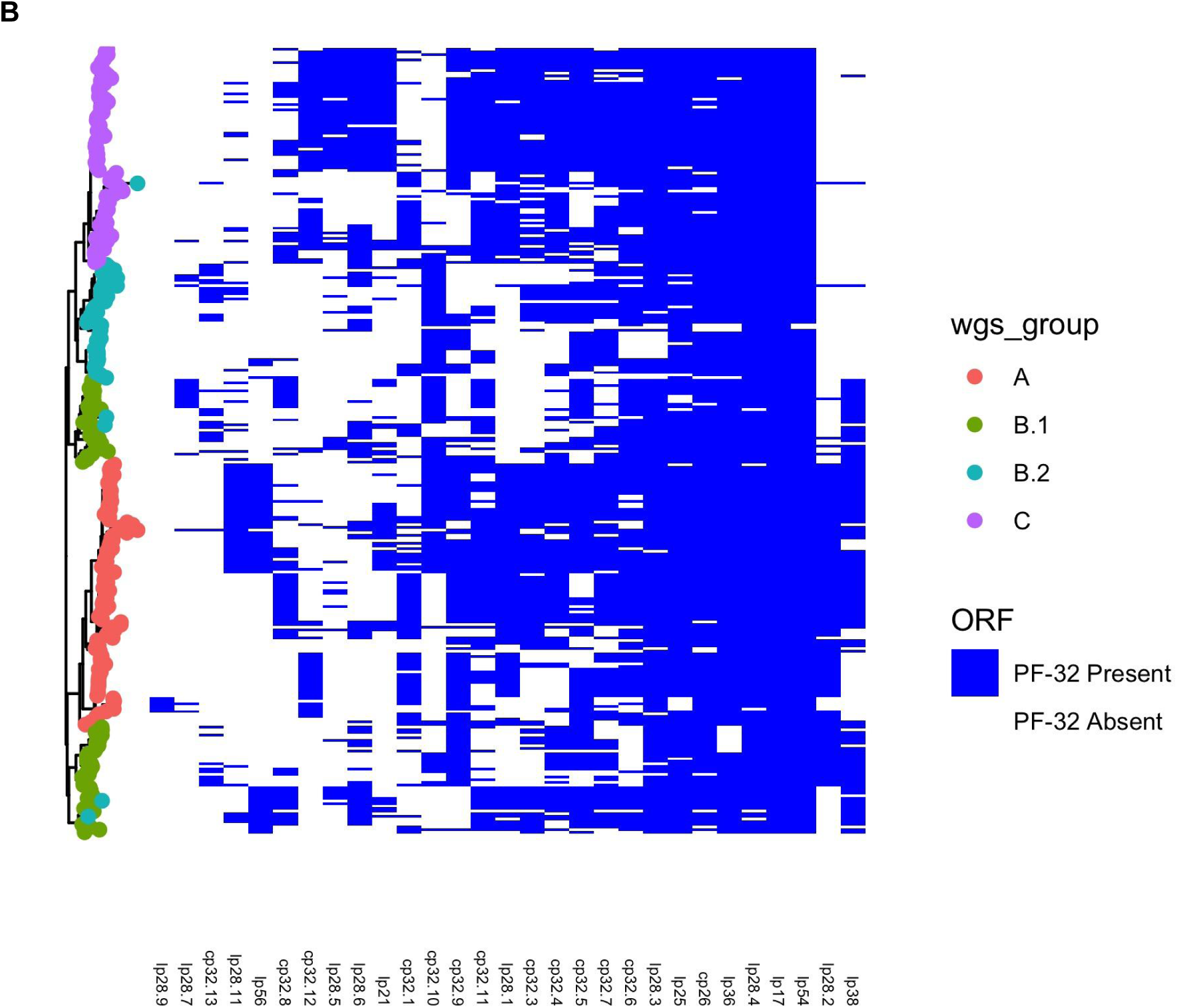
**A.** Phylogenetic tree created from the accessory genome with accessory genome elements plotted according to their presence/absence in individual strains. **B.** Phylogenetic tree created from the accessory genome with PFam32 plasmid compatibility sequences plotted according to the presence/absence in individual strains.

**Supplemental Figure 7:**
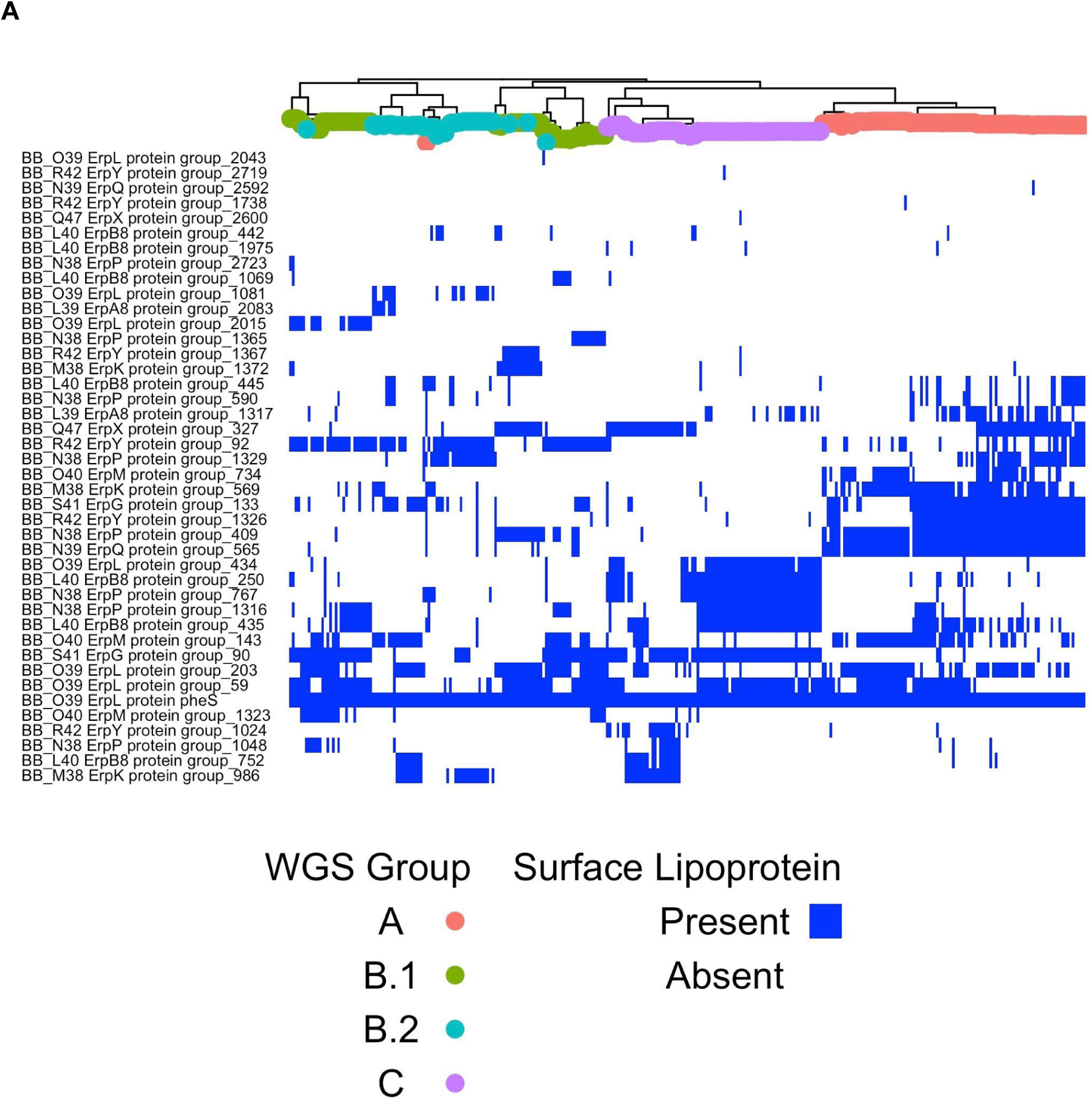
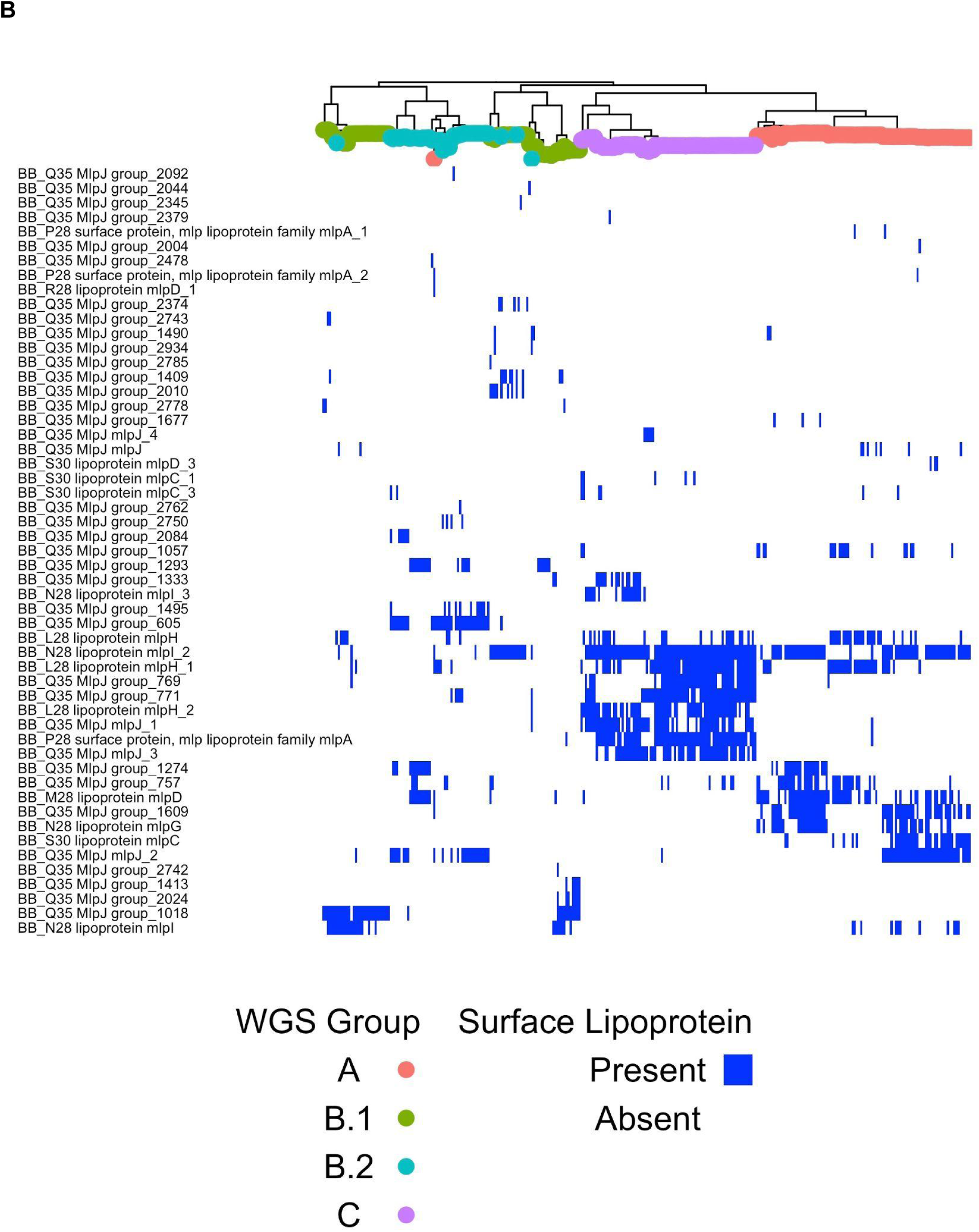
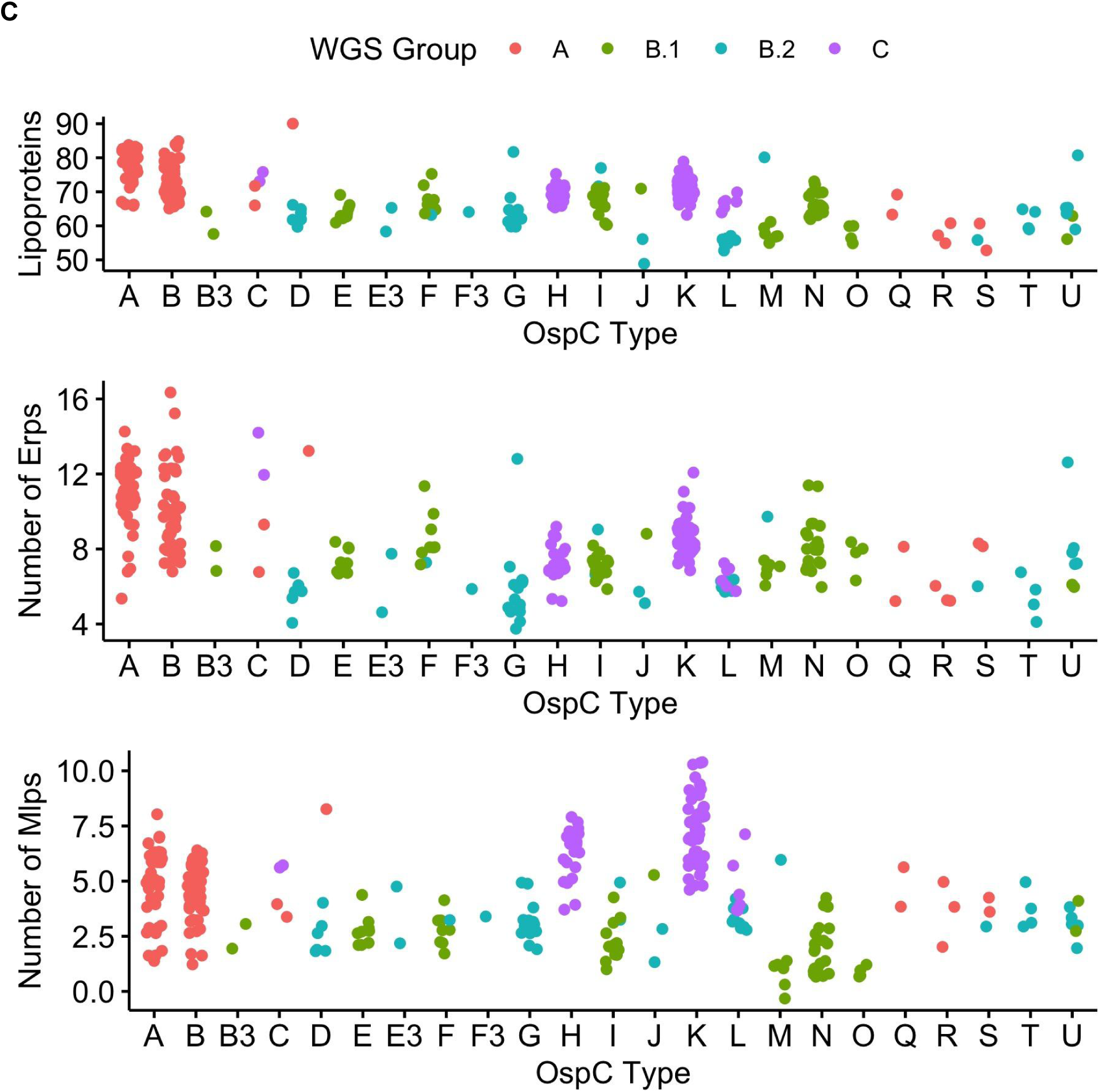
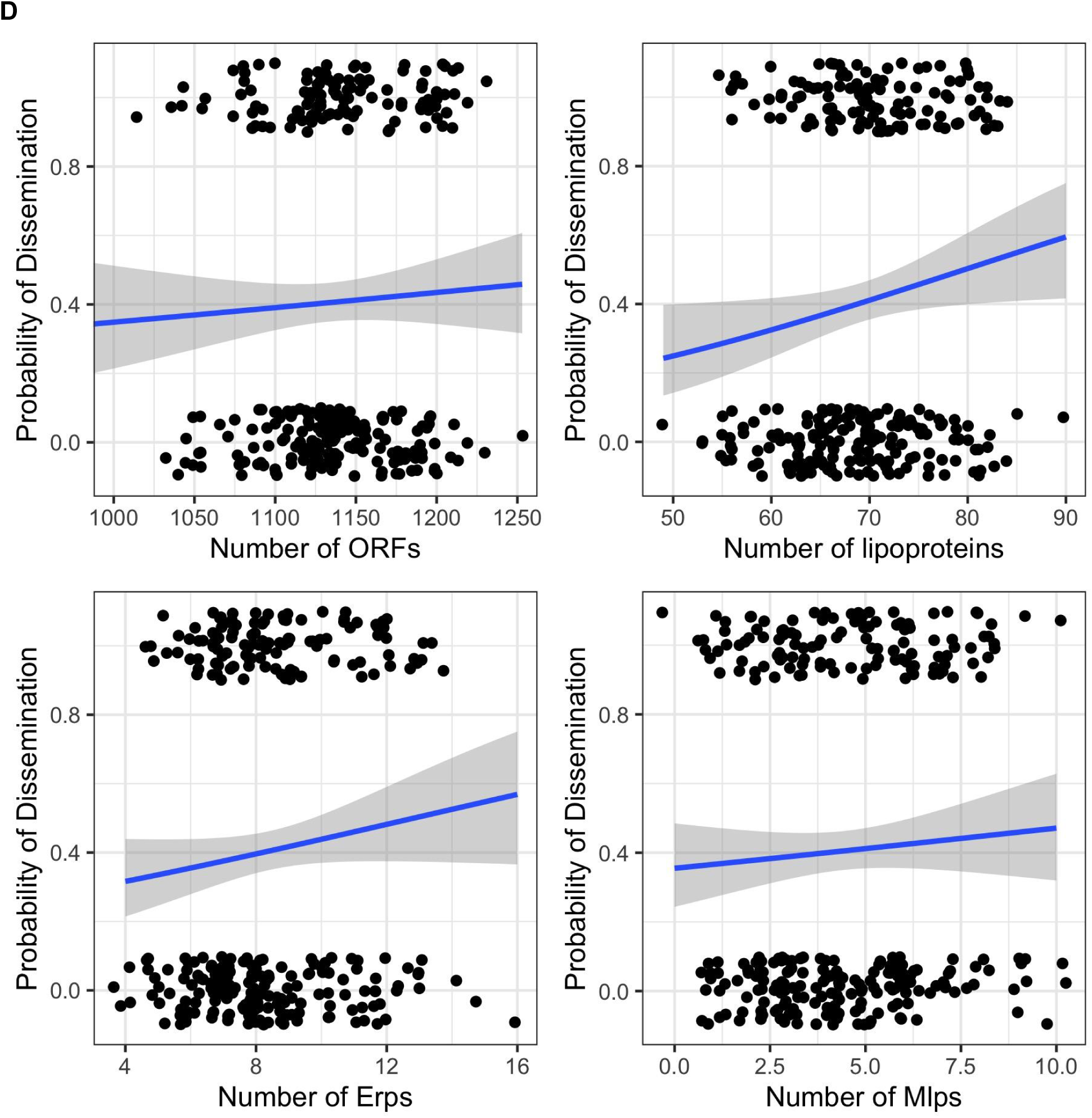
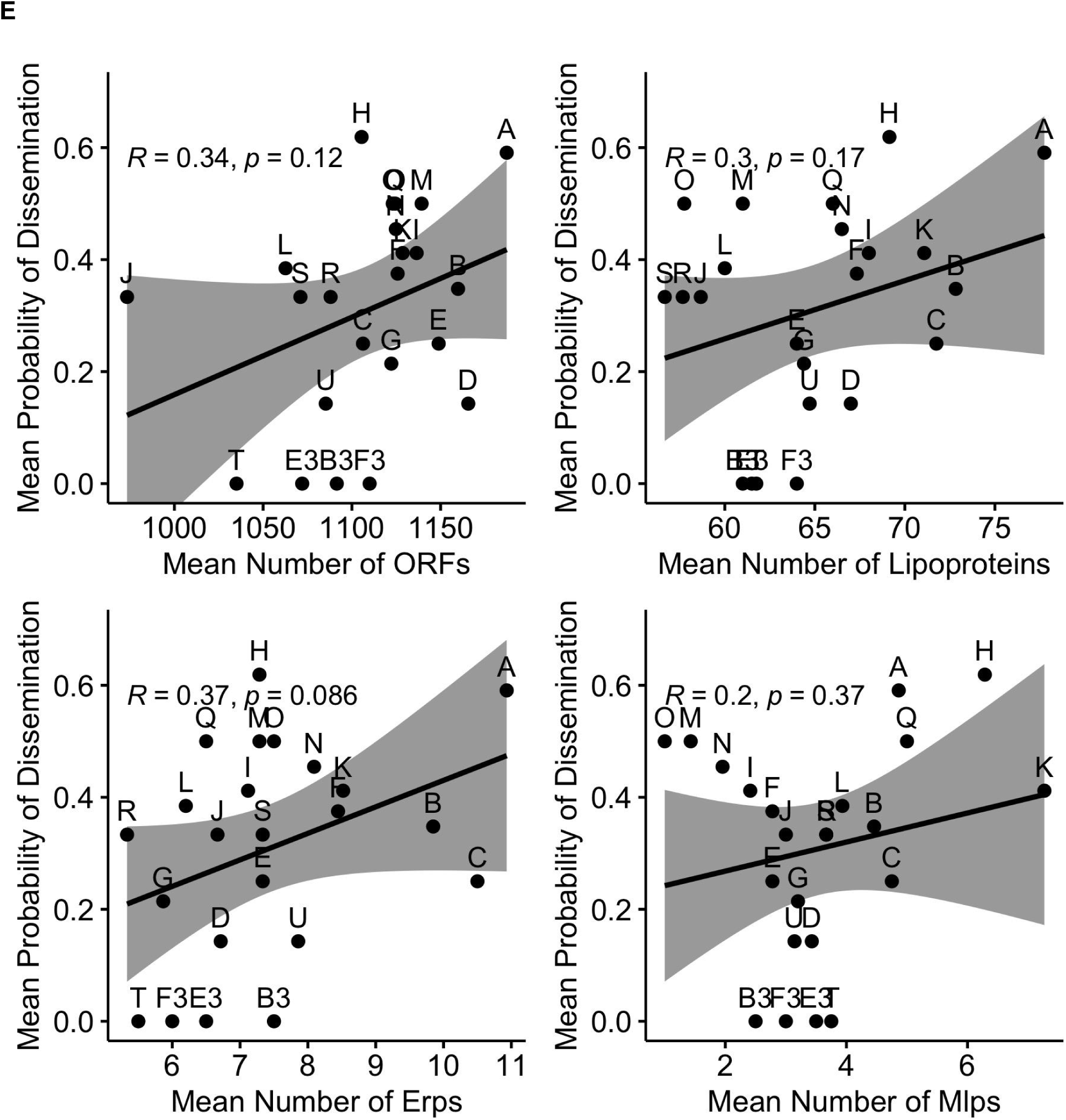
**A** and **B.** Core genome phylogeny with presence/absence of Erp (C) orthologs and Mlp (D) orthologs. **C.** The number of surface-exposed lipoproteins (top panel), Erps (middle panel), and Mlps (bottom panel) by OspC type. **D.** Probability of dissemination by number of ORF (top left, logistic regression coefficient for slope, β_1_ = 0.002 +/- 0.002, p = 0.450), number of surface-exposed lipoproteins (top right, β_1_ = 0.037 +/- 0.017, p = 0.03, logistic regression), number of Erps (bottom left, β_1_ = 0.087 +/- 0.053, p = 0.10, logistic regression), and number of Mlps (bottom right, β_1_ = 0.048 +/- 0.055 p = 0.38, logistic regression). **E.** For each OspC type, mean probability of dissemination vs mean number of ORF (top left), mean number of surface-exposed lipoproteins (top right), mean number of Erps (bottom left), and mean number of Mlps (bottom right).

**Supplemental Figure 8:**
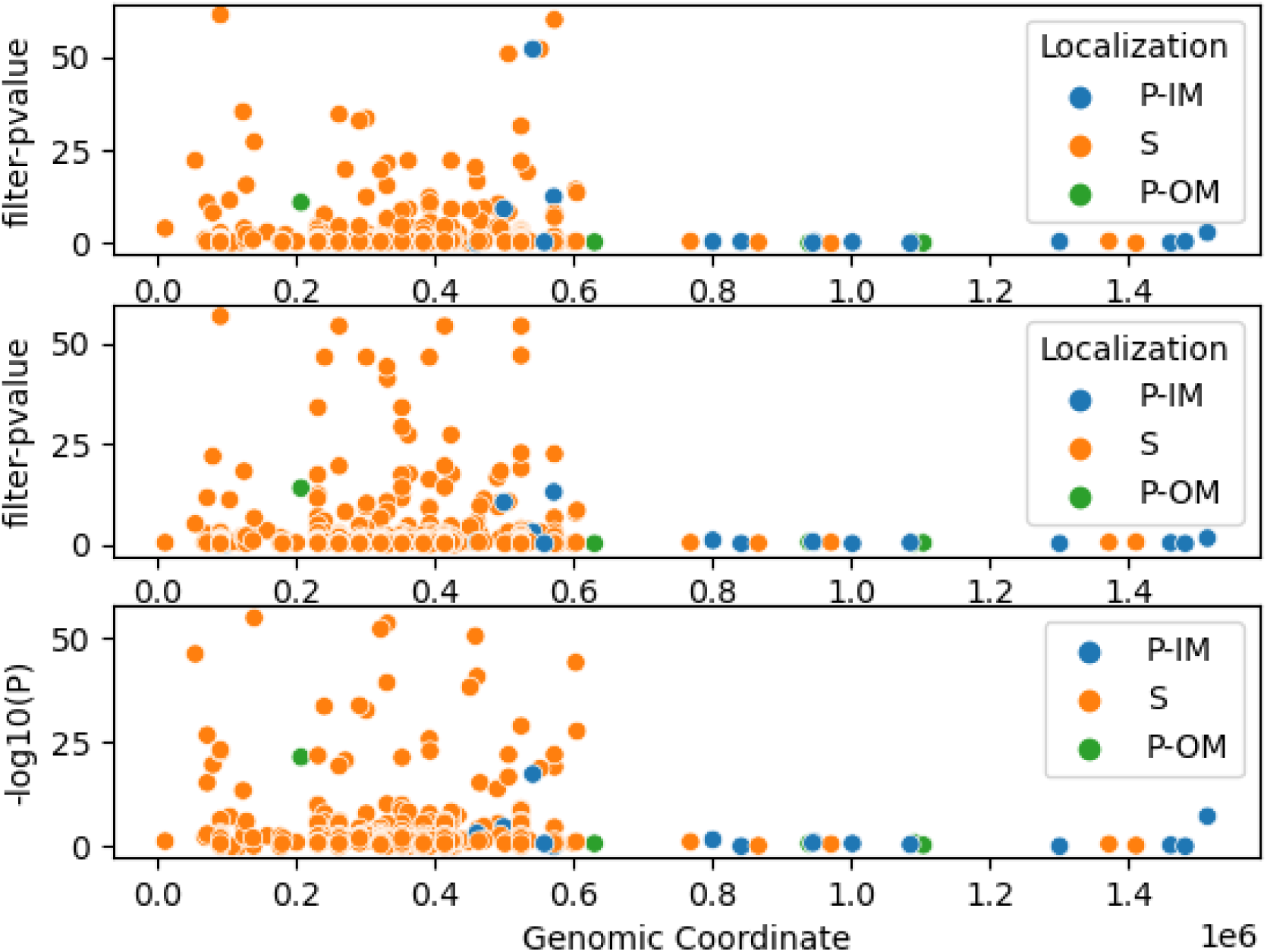
Manhattan Plots showing the association of individual lipoproteins with OspC type A (top panel), Osp C type K (middle panel), and RST1 (bottom panel). Individual lipoproteins are annotated by their localization. P-IM: Periplasmic inner membrane. P-OM: Periplasmic outer membrane. S: surface.

## List of supplemental data files

Supplemental Table 1: Summary table of isolates and phenotypes

Supplemental Table 2: List of isolates and phenotypes

Supplemental Table 3: Assembly statistics

Supplemental Table 4: Association statistics for plasmids, as inferred from PFam32 types.

Supplemental Table 5: Association statistics for plasmids, as inferred from B31 reference

Supplemental Table 5: Association statistics for lineage model

Supplemental Table 6: Association statistics for lineage model restricted to surface lipoproteins

Supplemental Table 7: Association statistics for OspC type A associations

Supplemental Table 8: Association statistics for OspC type K associations

Supplemental Table 9: Association statistics for RST1 associations

Supplemental Data File 1: List of ortholog groups with reference sequences

Supplemental Data File 2: High resolution version of presence/absence matrix in Figure 5B.

## Supplemental Note 1

The clock rate (in substitutions/site/year) for our initial model using a non-informative (CTMC rate reference) prior failed to converge—resulting in posterior 95% posterior density range from 5 x 10^-25^ substitutions/site/year to 1.2 x 10^-8^ substitutions/site/year—the implausibly small values at the lower end of the range are indicative of an insufficient temporal signal associated with genetic diversity in the core genome to establish an estimate without a priori assumptions. However, the inferred clock rate posterior had a clear single mode and a reasonable posterior mean (1.8 x 10^-9^ substitutions/site/year). To address this, we incorporated a priori information on mutation (gamma prior with shape 2, scale 1×10^-9^, for which 95% of the density is between 3.55 x 10^-10^ substitutions/site/year and 4.47 x 10^-9^ substitutions/site/year, concordant with previous suggestions that the rate is approximately 1 x 10^-9^ substitutions/site/year[52]). This analysis suggests that the common ancestry of circulating human-infectious populations was remote (95% posterior density for Midwest strains: 380,000 - 11.8 million years; 95% posterior density for Slovenian strains: 379,000 - 11.5 million years; all strains: 380,000 years, 11.8 million years) (Figure 2E-F). We also ran models with a fixed rate across a variety of reasonable values (1e-10 to 1e-8) (Figure 2G).

